# Membrane curvature-generating proteins crucial for autophagosome formation

**DOI:** 10.1101/2022.10.27.514035

**Authors:** Ning Wang, Yoko Shibata, Joao A. Paulo, Steven P. Gygi, Tom A. Rapoport

## Abstract

Autophagy is essential for cellular homeostasis and begins with the formation of a phagophore, a cup-like membrane sheet consisting of two closely apposed lipid bilayers connected by a highly curved rim. How the membrane sheet forms, bends, and eventually generates an autophagosome that enwraps cargo remains enigmatic. Specifically, it is unclear how the high membrane curvature of the phagophore rim, an energetically unfavorable state, is stabilized. Here, we demonstrate that phagophore formation requires the conserved, membrane curvature-generating REEP1 proteins. The REEP1 family proteins (REEP1-4 in vertebrates) differ from the related endoplasmic reticulum-shaping REEPs in abundance and membrane topology. In fission yeast, the single REEP1 ortholog is involved in both bulk and selective autophagy. It is recruited at early stages of phagophore formation and required for their maturation into autophagosomes. The function of REEP1 relies on its ability to generate high membrane curvature and its localization to the phagophore rim. Mammalian REEP1 proteins also associate with phagophores upon induction of autophagy and colocalize with early autophagic markers. We propose that REEP1 proteins stabilize the phagophore’s highly curved rim so that the two membrane sheets are kept in close proximity to form the autophagosome. Defective autophagy may underlie the effect of curvature-compromising mutations in human REEP1 proteins linked to neurological disease.

## Introduction

Macroautophagy (herein called autophagy) is induced by starvation, damage or overabundance of organelles, or the accumulation of misfolded proteins. Autophagy begins with the formation of a membrane sheet, called the phagophore or isolation membrane, which consists of two closely apposed phospholipid bilayers connected by a highly curved rim^1–4^ (**Extended Data Fig. 1a**). The membrane sheet expands and bends, generating a cup-like structure, and eventually closes on itself to form an autophagosome, in which the two membranes are no longer connected^5^. During closure, the autophagosome enwraps cytosolic components or organelles, either by random encapsulation (bulk autophagy) or by binding to specific receptors on the cargo (selective autophagy). The outer membrane of the autophagosome subsequently fuses with a lysosome (or vacuole in yeast), and both the inner membrane and the captured content are eventually degraded (**Extended Data Fig. 1a**).

Autophagy requires a number of conserved core components^1,6,7^ (**Extended Data Fig. 1a**). Some of these are involved in the attachment of a ubiquitin-like protein (Atg8 in yeast and LC3/GABARAP in mammals) to the lipid phosphatidylethanolamine in the phagophore membrane. Atg8 lipidation is critical for phagophore biogenesis and for completion of autophagy. However, how exactly the phagophore forms remains unclear. Specifically, it is unknown how the high curvature of the phagophore rim is stabilized. The rim has a radius of 7-15 nm in cross section^8^, which means that the phospholipid bilayer is extremely bent. Such high membrane curvature is energetically unfavorable and requires proteins for stabilization^9^. Some autophagy proteins such as Atg2 and Atg18 localize to the rim^10,11^, but none contain obvious domains that could stabilize such high curvature. The identification of phagophore rim-stabilizing proteins has been a long-standing question^8^, whose answer would provide important insight into how the unique structure of autophagosomes is generated.

In the case of the endoplasmic reticulum (ER), the high membrane curvature of tubules and sheet edges is generated by members of two distinct, evolutionarily conserved classes of proteins, the reticulons (Rtns) and a subfamily of the REEPs, that includes REEP5 in vertebrates and Yop1 in yeast^12–15^. The Rtns and REEP5/Yop1 proteins are abundant in most eukaryotic cells and contain four transmembrane segments (TMs) followed by an amphipathic helix (APH), all of which are required to generate high membrane curvature^16^. Although the Rtns and REEP5/Yop1 are the only known proteins that stabilize the edges of membrane sheets^17^, their absence in the budding yeast *Saccharomyces cerevisiae* does not affect autophagy^18^. These proteins are therefore unlikely to cause the membrane bending of the phagophore rim. However, the REEP1 proteins, a family related to the REEP5 proteins, could be involved, as they are much less abundant and therefore cannot play a major role in shaping the ER. REEP1 proteins are sequence-related to REEP5 proteins but lack the cytoplasmic N-terminus and first TM; the remaining three TMs are conserved and are predicted to form a similar structure as the last three TMs of REEP5 proteins (**Extended Data Fig. 1b** and **c**). REEP1 proteins are conserved in all eukaryotes, with the notable exception of budding yeasts (**Extended Data Fig 1d**). Here we show that the REEP1 proteins indeed play a crucial role in all forms of autophagy by stabilizing the highly curved rim of phagophores.

## Results

### REEP1 in fission yeast is important for all forms of autophagy

To investigate whether REEP1 proteins are required for autophagy, we took advantage of the fact that the fission yeast *Schizosaccharomyces pombe* possesses only a single ortholog of REEP1 (which we name Rop1, <u>R</u>eep <u>o</u>ne <u>p</u>rotein, encoded by gene *SPBC30D10*.*09c*). We first tested whether Rop1 participates in ERphagy, using Yop1 fused either to the fluorescent protein tdTomato (Yop1-tdT) or to GFP (Yop1-GFP) as specific markers. In logarithmically growing wild-type cells, Yop1-tdT and Yop1-GFP localized to the cortical ER as expected (**Fig. 1b** and **Extended Data Fig. 1e**). In contrast, they accumulated in vacuoles after ERphagy was induced, either by growing the cells to stationary phase, by nitrogen deprivation, or by triggering protein misfolding in the ER with dithiothreitol (DTT)^19^ (**Fig. 1b** and **Extended Data Fig. 1e**). Strikingly, no vacuolar accumulation of these ERphagy markers was seen in cells lacking Rop1 (*rop1*Δ) (**Fig. 1b** and **Extended Data Fig. 1e**), indicating a pronounced inhibition of autophagy. This defect was as strong as observed after deletion of Atg8 (*atg8*Δ). The only known ERphagy receptor in *S. pombe* is Epr1^7^, but its absence (*epr1*Δ) compromised autophagy only after DTT treatment (**Fig. 1b**).

**Fig. 1.**
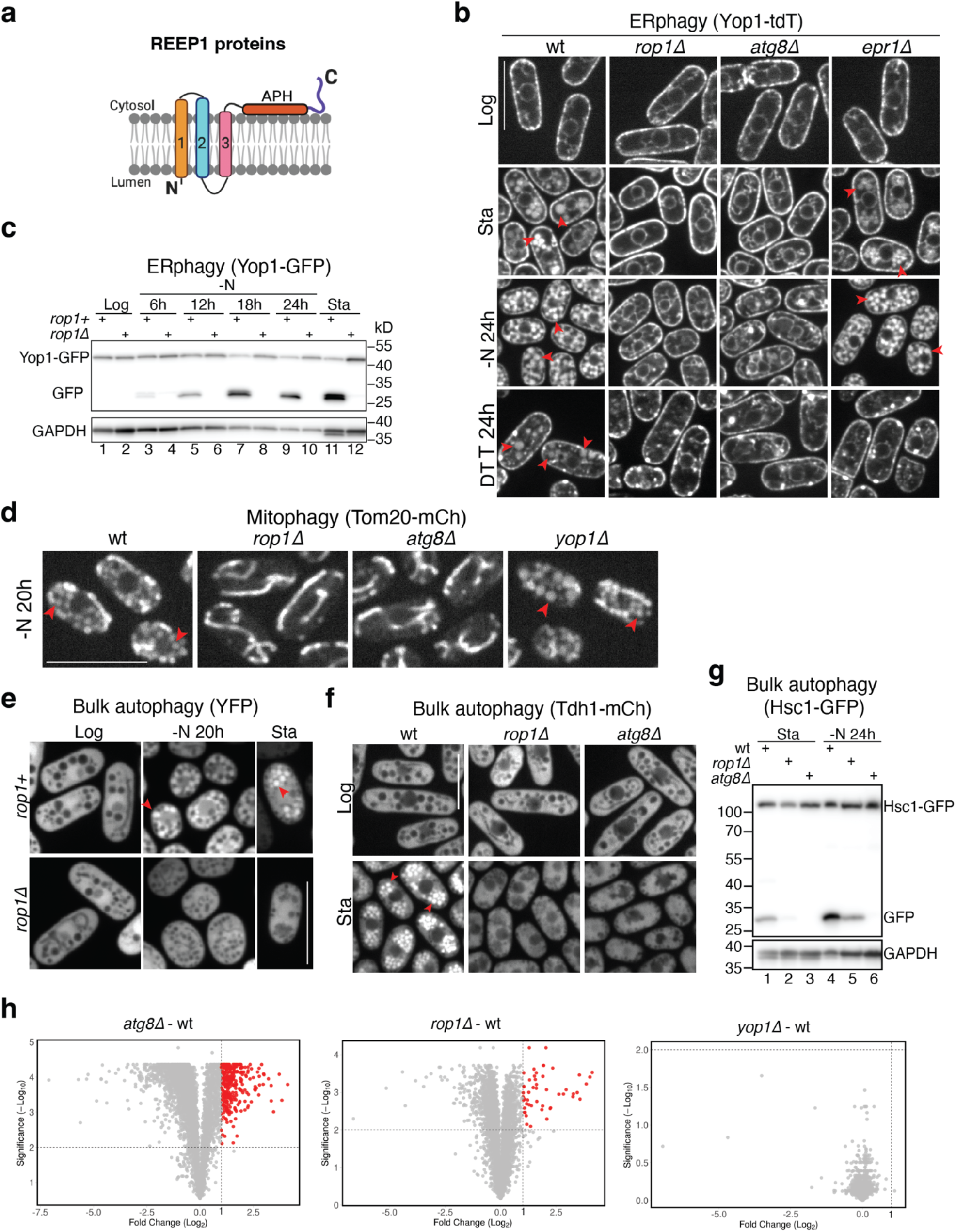
A crucial role for fission yeast Rop1 in autophagy. **a**, Membrane topology of REEP 1 proteins. Transmembrane (TM) segments are numbered. APH, amphipathic helix. **b**, The ER protein Yop1 was tagged with tdTomato (Yop1-tdT) at its genomic locus and expressed in wild-type (wt) *S. pombe* cells or cells lacking the indicated proteins. Cells in logarithmic (Log) or stationary (Sta) growth phase, cells nitrogen-starved for 24 h (-N 24h), or treated with DTT for 24 h (DTT 24h) were analyzed by fluorescence microscopy. Red arrowheads point to vacuoles. **c**, Yop1 was tagged with GFP (Yop1-GFP) and expressed in cells containing or lacking Rop1. Lysates from cells in Log or Sta phase, as well as from cells deprived of nitrogen (-N) for different time periods, were analyzed by SDS-PAGE and immunoblotting with GFP antibodies. Blotting with GAPDH antibodies served as loading control. **d**, As in b, but for the mitochondrial protein Tom20 tagged with mCherry (Tom20-mCh). **e**, As in b, but for the cytosolic protein YFP. **f**, As in b, but for the cytosolic protein Tdh1 tagged with mCherry (Tdh1-mCh). **g**, As in c, but for the cytosolic protein Hsc1 tagged with GFP (Hsc1-GFP). **h**, Protein accumulation in Rop1- or Atg8-lacking cells determined by quantitative isobaric tag-based proteomics. The abundance of proteins in *yop1Δ, rop1Δ*, and *atg8Δ* cells was compared with that in wt cells. Each point in the volcano plot represents a protein for which the ratio of its abundance in the mutant and in wt cells is given, as well as a measure of statistical significance (P-value), derived from a multiple non-parametric t-test based on three biological replicates. Proteins indicated by red points in the upper rectangle significantly accumulate in the mutant strain (P<0.01 and mutant-wt change > Log_2_ =1). Scale bars, 10 µm.

We further assessed ERphagy by monitoring the proteolytic cleavage of Yop1-GFP in vacuoles^19^. In cells grown to stationary phase or starved of nitrogen, the Yop1 portion of the fusion protein was digested while the GFP molecule resisted proteolysis (**Fig. 1c**). No processing of Yop1-GFP occurred in *rop1*Δ cells (**Fig. 1c**), consistent with a complete block of ERphagy. Cleavage of Yop1-GFP was also blocked in *rop1*Δ cells after inducing ERphagy with DTT or tunicamycin (**Extended Data Fig. 1f**). These defects were again as pronounced as observed in *atg8*Δ cells (**Extended Data Fig. 1f**). Similar results were obtained in stationary or nitrogen-starved cells using Yop1-mRFP, Rtn1-mRFP, or Rtn1-GFP as ERphagy markers (**Extended Data Fig. 1f** and **g**). Although Rop1 is homologous to Yop1 (**Extended Data Fig. 1b** and **c**), deletion of Yop1 (*yop1*Δ) had no effect on ERphagy (**Extended Data Fig. 1h**), as observed in *S. cerevisiae*^18^. The absence of the ER shaping protein Rtn1 (*rtn1*Δ) also had no effect (**Extended Data Fig. 1f**). Consistent with a crucial role for Rop1 in ERphagy, *rop1*Δ cells did not survive extended DTT treatment, similarly to *atg8*Δ or *epr1*Δ cells, whereas growth of wild-type and *yop1*Δ cells was unaffected (**Extended Data Fig. 1i**). The role of Rop1 in ERphagy is also conserved in the fission yeast *Schizosaccharomyces japonicus;* deletion of the single Rop1 homolog (*SJAG_05289*) prevented the accumulation of Rtn1 fused to mNeonGreen (Rtn1-mNG) inside vacuoles after inducing ERphagy with the Tor inhibitor rapamycin (**Extended Data Fig. 1j**). Taken together, these results establish that Rop1 is essential for ERphagy.

Rop1 is also critical for autophagy of mitochondria (mitophagy) and peroxisomes (pexophagy). A fusion of the mitochondrial membrane protein Tom20 with mCherry (Tom20-mCh) accumulated inside vacuoles in nitrogen-starved wild-type or *yop1*Δ cells, whereas this accumulation was greatly reduced in *rop1*Δ or *atg8*Δ cells (**Fig. 1d**). The absence of Rop1 also impaired the vacuolar cleavage of Tom20-mCh (**Extended Data Fig. 2a**). Likewise, both the vacuolar accumulation and proteolytic cleavage of the peroxisomal protein Pex11 fused to mCherry (Pex11-mCh) were reduced in *rop1*Δ cells (**Extended Data Fig. 2b** and **c**).

Finally, we found Rop1 to be important for bulk autophagy. The cytosolic marker proteins YFP, Tdh1-mCherry, and Sf3b-mCherry accumulated inside vacuoles of stationary or nitrogen-starved wild-type cells, but not in *rop1*Δ cells (**Fig. 1e** and **f**, and **Extended Data Fig. 2d**). Proteolytic cleavage of the cytosolic marker proteins Hsc1-GFP and Sec24-GFP, as well as of Pyk1-mCh and Pgk1-mCh, was also inhibited in *rop1*Δ cells under various autophagy-inducing conditions (**Fig. 1g**, and **Extended Data Fig. 2e** and **f**), although not as completely as in *atg8*Δ cells. We also tested the effect of Rop1 deletion on the turnover of Atg8 itself, as the Atg8 population bound to the inner autophagosomal membrane is normally degraded in vacuoles^3^ (**Extended Data Fig. 1a**). In wild-type cells, mEGFP-Atg8 indeed accumulated inside vacuoles upon starvation or DTT treatment (**Extended Data Fig. 2g** and **h**) and was proteolytically cleaved (**Extended Data Fig. 2i**). In *rop1Δ* cells, however, vacuolar accumulation and proteolytic cleavage were considerably reduced.

Quantitative isobaric tag-based proteomics showed that loss of Rop1 globally affected bulk autophagy. In starved cells, the absence of Rop1 caused the accumulation of various proteins (**Fig. 1h**), most of which are cytosolic, whereas no significant changes occurred in the absence of Yop1 (**Fig. 1h**). A large number of proteins was also stabilized in cells lacking Atg8, as expected. Most of the significantly accumulating proteins detected in *rop1*Δ overlapped with those in *atg8*Δ cells (**Extended Data Fig. 3a**), consistent with Rop1 and Atg8 acting in the same pathway.

Rop1’s role in autophagy is also supported by thin-section electron microscopy (EM) (**Extended Data Fig. 3b-d**). In nitrogen-starved wild-type or *yop1Δ* cells, many vacuoles contained electron-dense material that likely corresponds to residual internal membranes of autophagosomes (**Extended Data Fig. 3b** and **c**). In contrast, vacuoles were empty in *rop1Δ* and in *atg8Δ* cells (**Extended Data Fig. 3b** and **c**). The number of vacuoles in *rop1Δ* cells was also more variable than in wild-type cells (**Extended Data Fig. 3c**) and many were smaller (yellow arrows in **Extended Data Fig. 3b**). Similar results were obtained with cells in stationary phase (**Extended Data Fig. 3d**). Light microscopy showed that the loss of Rop1 did not affect ER morphology or the localization of components of the secretory pathway (**Extended Data Fig. 3e**), except that the vacuoles were smaller in starved cells (**Extended Data Fig. 3f**), likely caused by defective autophagy.

Although Rop1 has a similar membrane topology as Atg40 in *S. cerevisiae*, it is not a functional homolog. Atg40 serves as a cargo receptor for ERphagy, but is not found outside budding yeasts^20,21^. Atg40 contains a typical Atg8-interacting motif (AIM) that binds Atg8 *in vitro* and is required for the receptor’s function in ERphagy^20,21^ (**Extended Data Fig. 4c**). In contrast, although Rop1 contains a potential AIM in its C-terminal segment, ablation of this motif had no effect on autophagy (**Extended Fig. 4a** and **b**), and neither wild-type Rop1 nor an AIM mutant bound Atg8 *in vitro* (**Extended Data Fig. 4c**). *S. cerevisiae* Atg40 also failed to complement the growth defect of the *S. pombe rop1Δ* mutant on DTT (**Extended Data Fig. 4d**). Taken together, our results indicate that Rop1 is fundamentally involved in all forms of autophagy and is not simply an ERphagy receptor.

### Crucial role of Rop1 in phagophore expansion and autophagosome formation

To understand the function of Rop1 in autophagy, we first compared the localization of various autophagy factors in wild-type versus *rop1Δ* cells under starvation conditions. Using Atg8 fused to mEGFP (mEGFP-Atg8) as a marker of autophagosomes, we noted that most wild-type cells contained two or three mEGFP-Atg8 punctae (**Fig. 2a** and **b**) that likely represent nascent phagophores. In the absence of Rop1, the number of punctae increased while their brightness decreased (**Fig. 2a** and **b**), as previously observed for *atg2Δ* and *atg18Δ* mutants^22^. In wild-type cells, the autophagy components Atg1 (the earliest autophagy marker^23^; see **Extended Data Fig. 1a**), Atg9 (a multi-spanning membrane protein of autophagosomes)^24^, Atg5 (a component of the ubiquitin-like conjugation machinery)^25^, and Atg2 (probably a phospholipid transfer protein)^26^ all colocalized with mEGFP-Atg8 (**Fig. 2c-f**). In contrast, most of the mEGFP-Atg8 punctae in *rop1Δ* cells still contained Atg1 (**Fig. 2c**) but showed reduced recruitment of Atg9, Atg5, and Atg2 (**Fig. 2d-f**). Thus, the earliest autophagic structures can still form in the absence of Rop1, but phagophore expansion and autophagosome formation are delayed or stalled.

**Fig. 2.**
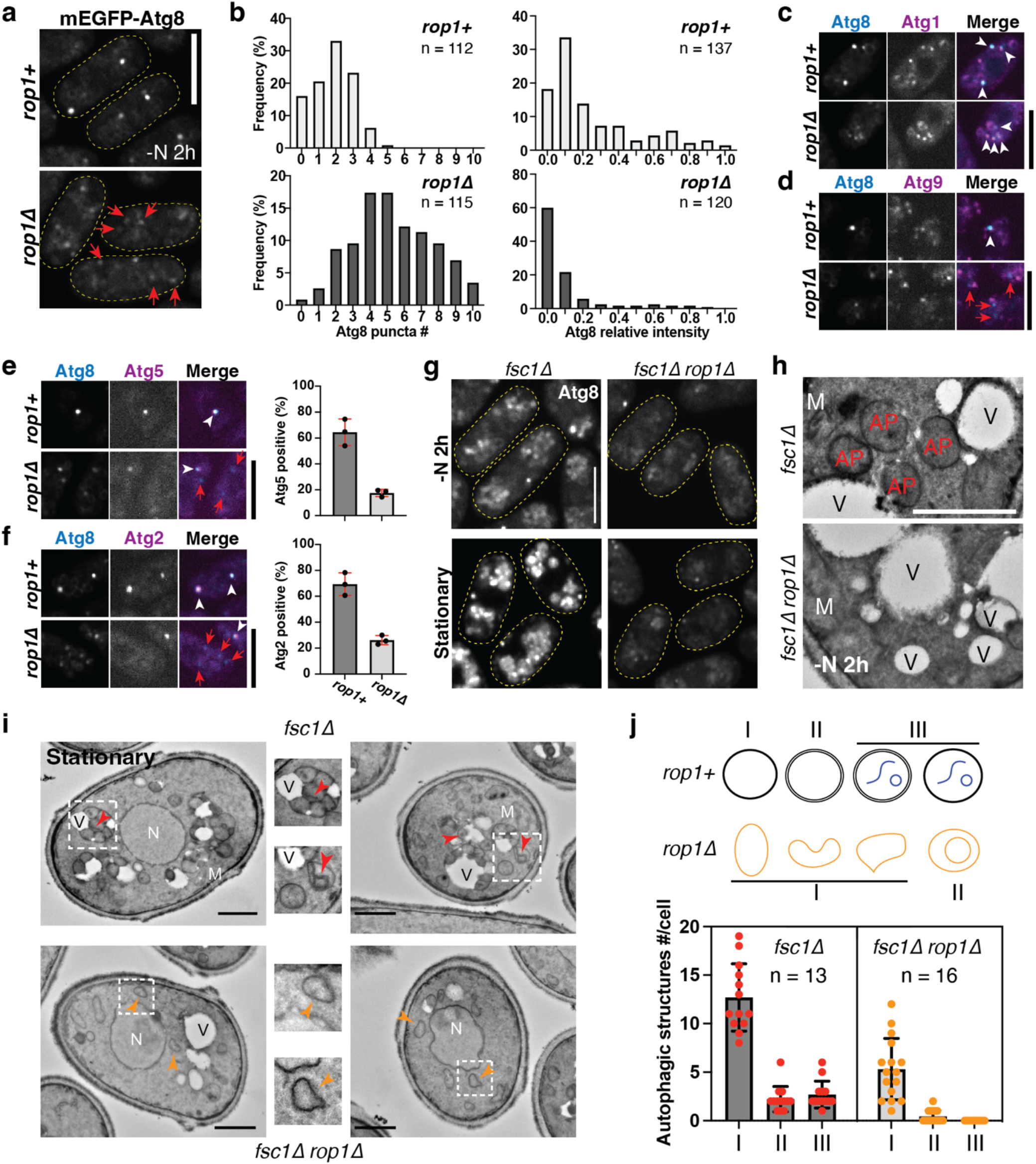
Rop1 is required for autophagosome formation. **a**, Atg8 was tagged with mEGFP (mEGFP-Atg8) and expressed at its native level in *rop1*^*+*^ or *rop1Δ* cells. Cells were imaged after 2h of nitrogen starvation (-N 2h). The boundaries of cells are indicated by dashed lines. Dim Atg8 punctae in *rop1Δ* cells are indicated by red arrows. Scale bar, 5 µm. **b**, Shown are the number of mEGFP-Atg8 punctae per cell (left) and their intensity, expressed relative to the brightest Atg8 punctum in *rop1*^*+*^ cells (right). n, number of analyzed cells (left) or punctae (right). **c**, Atg1 was tagged with tdTomato and visualized together with mEGFP-Atg8 in *rop1*^*+*^ or *rop1Δ* cells after 2h of nitrogen starvation. Scale bar, 5 µm. **d**, As in c, but with tagged Atg9. **e**, As in c, but with tagged Atg5. The right panel shows the percentage of Atg8 punctae containing Atg5 (mean and standard deviation). **f**, As in e, but with tagged Atg2. **g**, As in a, but with *fsc1Δ* or *fsc1Δ rop1Δ* cells. The cells were starved for 2h (-N 2h) or in stationary (Sta) growth phase. Scale bar, 5 µm. **h**, *fsc1Δ* or *fsc1Δ rop1Δ* cells were starved for 2h (-N 2h) and analyzed by TEM. M, mitochondrion; V, vacuole; AP, autophagosome. **i**, As in h, but for stationary cells. N, nucleus. Red arrowheads indicate autophagosomes with enclosed substrate and orange arrowheads point to aberrant autophagic structures. The regions in the squares are magnified in the central images. Scale bar, 1 µm. **j**, Quantification of autophagic structures in i. The scheme shows the structures that were counted. The bars show means and standard deviations. *n*, number of cells analyzed.

To further test whether Rop1 affects autophagosome formation, we prevented autophagosome fusion with the vacuole by deleting the fusion factor Fsc1^22^. After nitrogen starvation or in stationary phase, the number of mEGFP-Atg8 punctae increased in cells lacking Fsc1 (*fsc1Δ*) (**Fig. 2g**), as reported previously^22^. This accumulation was almost entirely abolished in *fsc1*Δ cells devoid of Rop1 (*fsc1*Δ *rop1Δ*). By thin-section EM, numerous spherical structures tethered to vacuoles were apparent in nitrogen-starved or stationary *fsc1Δ* cells (**Fig. 2h-j**). These structures correspond to autophagosomes because they contain engulfed material (red arrows in **Fig. 2i**) and are surrounded by two closely spaced membranes (**Fig. 2i**). In contrast, the number of autophagosomes was significantly reduced in *fsc1*Δ *rop1Δ* cells, and the remaining ones were irregularly shaped (orange arrows in **Fig. 2i**; quantification in **Fig. 2j**). Although some still showed two bilayers, the membranes were much more separated than in *fsc1Δ* cells and the structures did not contain encapsulated cargo (quantification in **Fig. 2j**). Similar results were obtained after DTT treatment (**Extended Data Fig. 4e-g**). Rop1 is thus required for proper autophagosome formation.

### Generation of membrane curvature underlies Rop1’s function in autophagy

Although REEP1 subfamily members have one less TM than the ER-shaping REEP5 homologs (**Extended Data Fig. 1b**), they also generate high membrane curvature. When *S. japonicus* Rop1 (sjRop1) was expressed in bacteria, a significant fraction of the protein did not sediment with membranes (**Fig. 3a**). Particles purified from this soluble fraction had diameters of ∼10 nm when viewed by negative-stain EM (**Fig. 3b**). Such a small diameter is not theoretically possible for vesicles with a lipid bilayer^16^. Thus, sjRop1 generates high membrane curvature like Yop1 and other REEP5 homologs^16^, causing the conversion of a bilayer into a monolayer to form a lipoprotein particle. Generation of high curvature was also observed after sjRop1 was solubilized from the membrane fraction using detergent (**Extended Data Fig. 5a**): upon reconstitution with phospholipids, detergent-purified sjRop1 readily formed lipoprotein particles or very small vesicles (**Fig. 3c**). The APH in sjRop1 (**Fig. 1a**) is required for curvature generation, as its deletion (Δ107-120) resulted in large low-curvature vesicles whereas a more C-terminal truncation (Δ126-170) had no effect (**Fig. 3c**; for purity of the proteins see **Extended Data Fig. 5b**). In addition, sjRop1 forms homodimers just like REEP5/Yop1^16^, as shown by photo-crosslinking with probes incorporated into the first TM (**Extended Data Fig. 5d**). This TM is predicted to reside at the interface between the two monomers and is equivalent to the second TM of REEP5/Yop1 proteins (**Extended Data Fig. 5c**).

**Fig. 3.**
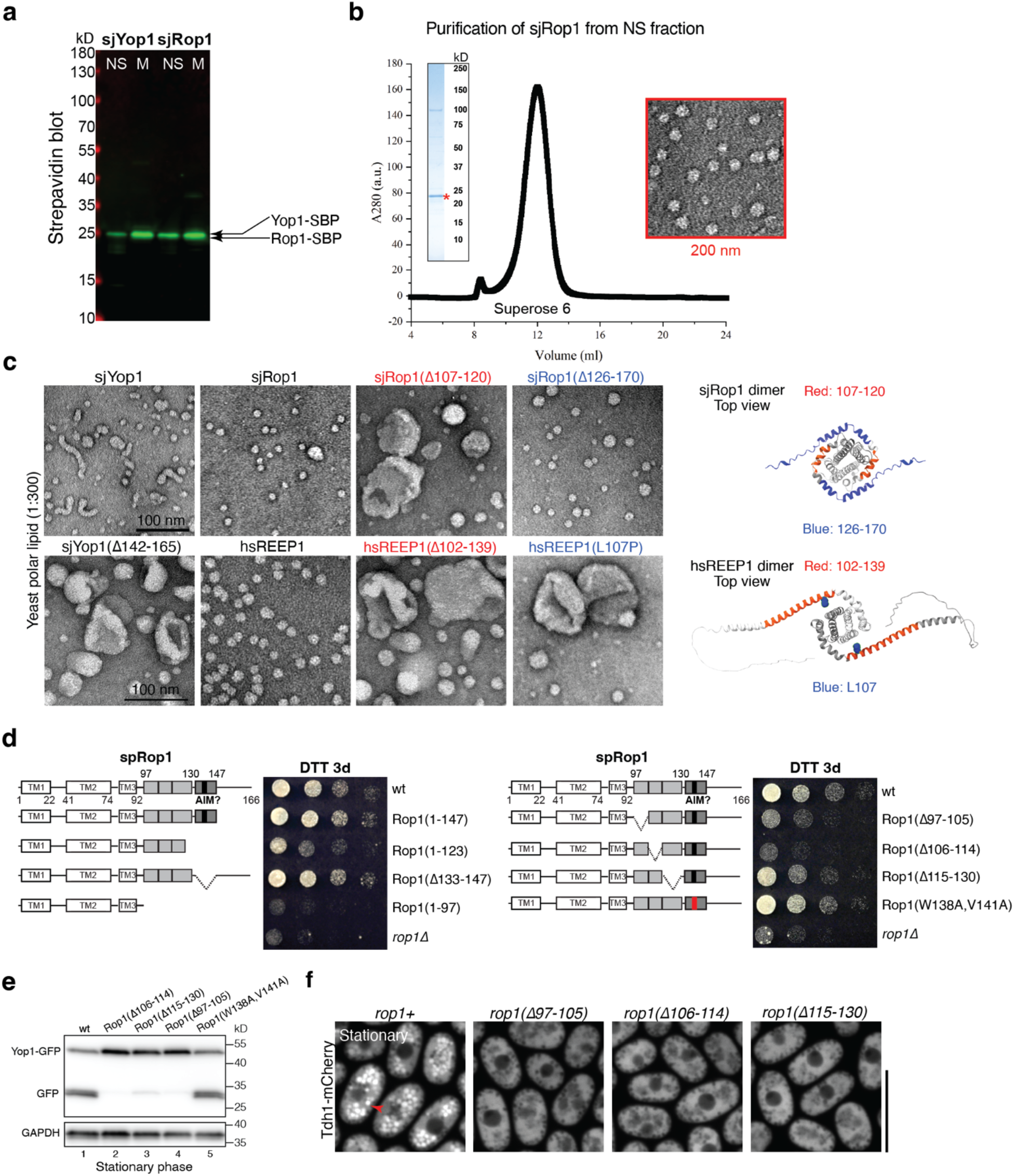
Rop1’s function in autophagy requires membrane curvature generation. **a**, SBP-tagged sjRop1 and sjYop1 from *S. japonicus* were expressed in *E. coli*. Cell lysates were subjected to ultracentrifugation and the membrane (M) and non-sedimentable (NS) fractions analyzed by SDS-PAGE, followed by blotting with fluorescently labeled streptavidin. **b**, sjRop1 of the NS fraction was bound to streptavidin beads, the tag was removed with TEV protease, and the eluted material subjected to gel filtration. The peak fraction was analyzed by Coomassie-blue staining (left inset) and negative-stain EM (right inset; square size 200 nm). **c**, sjYop1, sjRop1, hsREEP1 or the indicated mutants were purified from the membrane fraction and mixed with liposomes. The detergent was removed, and the reconstituted material was analyzed by negative-stain EM. Scale bars, 100 nm. The panels on the right show topdown views of the Alphafold-predicted protein structures, with the regions deleted in the mutants and position 107 of hsREEP1 highlighted. **d**, Wild-type (wt) *S. pombe* Rop1 (spRop1) or the indicated mutants were expressed from the endogenous genomic locus. The cells were treated with DTT for three days (DTT 3d) and plated after serial dilution. The schemes show the domains of spRop1, with the APHs and putative AIM motif in grey and black, respectively, and deleted regions indicated. The W138A, V141A mutant is designed to abrogate the AIM motif. **e**, The ER protein Yop1 was tagged with GFP (Yop1-GFP) and expressed in wt cells or cells expressing the indicated Rop1 mutants. Cell lysates were analyzed by SDS-PAGE and immunoblotting with GFP antibodies. Blotting with GAPDH antibodies served as loading control. **f**, The cytosolic protein Tdh1 was tagged with mCherry (Tdh1-mCh) at its genomic locus and expressed in wt cells or cells expressing the indicated Rop1 mutants. Stationary phase cells were analyzed by fluorescence microscopy. Scale bar, 10 µm.

Purified human REEP1 (hsREEP1) behaved analogously to sjRop1 (**Extended Data Fig. 5e**). Again, mutations in the APH (Δ102-139 and L107P) compromised REEP1’s ability to generate lipoprotein particles or small liposomes (**Fig. 3c**). Notably, these mutations are associated with the human diseases distal hereditary motor neuropathy type V (HMN5B)^27^ and hereditary spastic paraplegia (HSP)^28^, respectively.

Finally, we confirmed that Rop1’s ability to generate high membrane curvature is required for autophagy. Even small deletions in the APH reduced cell survival on DTT (**Fig. 3d**) and vacuolar cleavage of Yop1-GFP (**Fig. 3e**), and also prevented the vacuolar accumulation of cytosolic Tdh1-mCh (**Fig. 3f** and **Extended Data Fig. 5f**).

### Rop1 localizes to the phagophore rim

The ability of Rop1 to generate high membrane curvature raised the possibility that the protein might localize to the phagophore rim. To test this idea, we first examined the subcellular localization pattern of Rop1. Regardless of whether autophagy was induced or not, Rop1 fused to mNeonGreen (Rop1-mNG) localized to numerous punctae (**Fig. 4a** and **b**, and **Extended Data Fig. 6b**), which are likely vesicles. When overexpressed, Rop1-mNG moved to the cortical ER and nuclear envelope and colocalized with Yop1 (**Extended Data Fig. 6c**), suggesting that the vesicles originate from the ER. Following induction of autophagy by nitrogen starvation, Rop1-mNG became strongly enriched on ≥80% of phagophores marked by mCherry-Atg8 (mCh-Atg8) (**Fig. 4a**, quantification in **Fig. 4b**). Live-cell imaging revealed that Rop1-mNG and mCh-Atg8 were simultaneously recruited to phagophores, suggesting early Rop1 recruitment to the forming autophagosome (**Extended Data Fig. 6a**). Similarly, Rop1-mNG colocalized with the majority of phagophores marked by Atg2-tdTomato (Atg2-tdT) (**Fig. 4a-c**). These data show that a fraction of the cellular pool of Rop1 moves to phagophores after autophagy is induced.

**Fig. 4.**
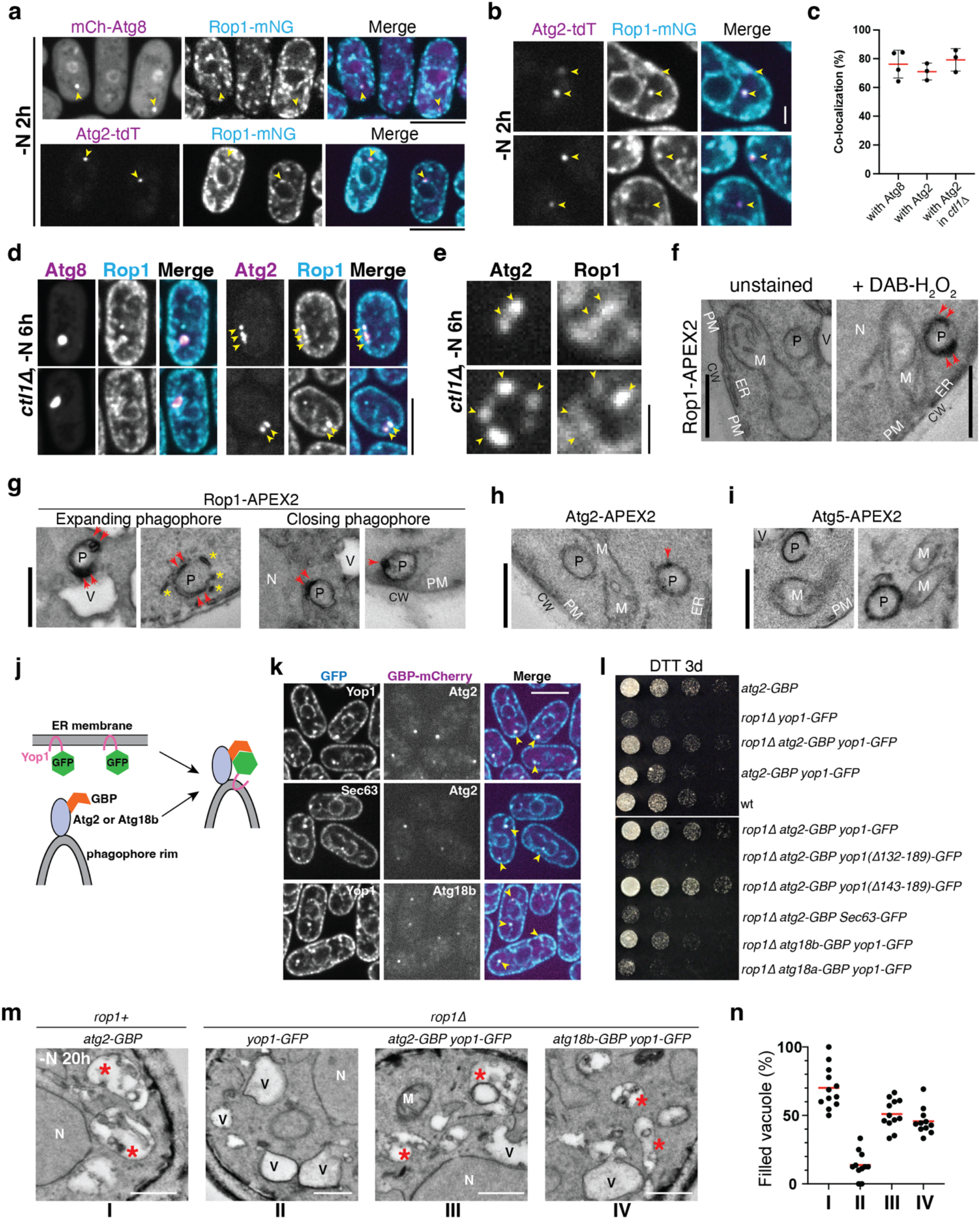
Rop1’s function requires localization to the curved phagophore rim. **a**, Rop1 was tagged with mNeonGreen (Rop1-mNG) and expressed together with mCherry tagged Atg8 (mCh-Atg8) or tdTomato tagged Atg2 (Atg2-tdT). Cells were imaged after 2h of nitrogen starvation. Arrowheads point to punctae where colocalization is observed. Scale bar, 5 µm. **b**, As in a, showing additional examples of colocalization of Rop1-mNG and Atg2-tdT. Scale bar, 1 µm. **c**, Quantification of co-localization of Rop1 with Atg8 and Atg2 in starved wild-type and *ctl1Δ* cells. n = 50-100 punctae from 3 or 4 independent experiments. Shown are the means and standard deviations. **d**, As in a, but with *ctl1Δ* cells. Arrowheads point to colocalization of Rop1 with Atg2 on enlarged phagophores. Scale bar, 5 µm. **e**, As in d, showing additional examples of colocalization of Rop1-mNG and Atg2-tdT in a magnified view. Scale bar, 1 µm. **f**, Cells expressing Rop1-APEX2 were nitrogen-starved for 2h, fixed, stained with diaminobenzidine (DAB) and H_2_O_2_, and analyzed by TEM. A fixed sample without DAB-H_2_O_2_ treatment (unstained) is shown as a control. P, phagophore; CW, cell wall; M, mitochondrion; N, nucleus; PM, plasma membrane; V, vacuole; ER, endoplasmic reticulum. Arrowheads point to the tips of a C-shaped phagophore. Scale bar, 500 nm. **g**, As in f, except that the closing phagophores are shown on the right. Asterisks point to small vesicles or short tubules near the expanding phagophore. **h**, As in f, but with Atg2-APEX2. **i**, As in f, but with Atg5-APEX2. **j**, Scheme showing how a fusion of Atg2 with a GFP nanobody (Atg2-GBP) drags Yop1-GFP from the ER to the phagophore rim. A similar strategy was used with Atg18b-GBP. **k**, Atg2-GBP or Atg18b-GBP were tagged with mCherry and expressed together with Yop1-GFP or Sec63-GFP. The cells were imaged after 2h of nitrogen starvation. Arrowheads point to colocalization of the coexpressed proteins. **l**, Wild-type (wt) *S. pombe* cells or the indicated mutants were treated with DTT for three days (DTT 3d) and plated after serial dilution. **m**, Wt or *rop1Δ* cells expressing the indicated fusion proteins were starved for 20h and analyzed by TEM. Asterisks point to vacuoles containing electron-dense material, likely residual membranes of autophagosomes. V, empty vacuole. M, mitochondrion. N, nucleus. Scale bars, 500 nm. **n**, Quantification of the results in m. Shown is the percentage of filled vacuoles per cell. The red lines show the mean.

Next, we asked whether the phagosomal localization of Rop1 corresponds to the rim. To facilitate detection of the rim, we used a mutant, *ctl1Δ*, in which phagophores are abnormally enlarged^22^. mCh-Atg8 localized in *ctl1*Δ cells on bright structures that were often cup-shaped (**Fig. 4d**), as expected. Importantly, Rop1 localized only to subregions of these structures (**Fig. 4d**), which were also enriched for Atg2 (**Fig. 4d** and **e**, quantification in **Fig. 4c**), a protein well-known to localize to the phagophore rim in multiple organisms including *S. pombe*^10,11,22,26^. These results thus suggest that Rop1 concentrates at the rim of expanding phagophores.

To support this conclusion, we fused Rop1 to ascorbate peroxidase 2 (Rop1-APEX2) and examined the localization of the fusion protein by thin-section EM. Rop1-APEX2 was visualized as dark regions that were particularly intense at the tips of cup-shaped expanding phagophores (**Fig. 4f** and **g**). Small structures near phagophores, likely Rop1-containing vesicles, were also labeled (**Fig. 4g**). Rop1-APEX2 also localized to one or two closely spaced dots on spherical structures that likely correspond to phagophores about to close and become autophagosomes (**Fig. 4g**). Atg2 showed a similarly restricted localization on these structures (**Fig. 4h**). In contrast, Atg5 was evenly distributed along them (**Fig. 4i**), consistent with the known localization of this factor over the entire phagophore^10,22^. These data support the restricted localization of Rop1 to the phagophore rim, analogously to Atg2. Rop1-APEX2 was also seen at the cortical ER, and in vesicles close to the ER and nuclear envelope (**Extended Data Fig. 6d**), consistent with the localization of this protein by light microscopy (**Extended Data Fig. 6b**).

Finally, we tested whether Rop1’s localization to the phagophore rim is required for autophagy. We harnessed Atg2 to recruit Yop1 (which also generates high membrane curvature but does not normally participate in autophagy, see above) to the phagophore rim, and asked whether it could rescue the autophagic defects that result from the absence of Rop1. Relocalization of Yop1 was achieved by expressing Yop1-GFP together with Atg2 fused to a GFP-binding protein (GBP, a GFP nanobody)^29^ (**Fig. 4j**). A sub-population of Yop1-GFP indeed moved to phagophores upon starvation, as shown by colocalization with Atg2-GBP-mCherry (**Fig. 4k**). Importantly, rim-localized Yop1 completely rescued the growth defect of *rop1Δ* cells on DTT (**Fig. 4l**) and increased the number of filled vacuoles in these cells to wild-type levels (**Fig. 4m** and **n**). Rescue required Yop1’s ability to generate high membrane curvature, as relocalization of Yop1-GFP lacking the APH, Δ132-189, failed to restore growth, in contrast to a more distal truncation, Δ143-189, that retains the APH (**Fig. 4l**, and **Extended Data Fig. 6e**). Growth also failed to be restored when the bulk ER protein Sec63 was recruited similarly to the phagophore rim (**Fig. 4k** and **l**). On the other hand, recruitment of Yop1 to the phagophore rim by Atg18b^22^, a known interacting partner of Atg2, rescued autophagy in *rop1Δ* cells, although the rescue was less pronounced (**Fig. 4k-n**). The homolog Atg18a was ineffective in this assay (**Fig. 4l** and **Extended Data Fig. 6e**), consistent with reports that it differs from Atg18b in localization and function^22^. Taken together, these data demonstrate that the function of Rop1 in autophagy correlates with the protein’s localization to the phagophore rim.

### Mammalian REEP1 proteins localize to phagophores

Lastly, we tested whether mammalian REEP1 proteins also localize to autophagic structures. Without induction of autophagy, all REEP1 homologs localized prominently to vesicles and short tubules in human U2OS and SKN-SH cells (**Extended Data Fig. 7a** and **b**, and **d-j**). Most molecules stayed in these vesicles even after autophagy was induced, as shown for stably expressed REEP1 fused to mEmerald (REEP1-mEm) (**Extended Data Fig. 7i**). However, a fraction became enriched on punctae formed by the Atg8 orthologs LC3 (**Fig. 5a**) and GABARAP (**Extended Data Fig. 8a**). Notably, these REEP1-LC3 punctae lacked the lysosomal marker LAMP1, indicating that they correspond to phagophores or autophagosomes not yet fused with lysosomes (**Fig. 5a**). A fraction of REEP1 also colocalized with various autophagic receptors and cargo, which included the ERphagy receptor FAM134B^30^; the cytosolic cargo receptor p62/SQSTM1^31^; and cytosolic aggregates of a GFP-fusion of mutant superoxide dismutase 1 (SOD1 (A4V)-GFP)^32^, a disease protein in familial amyotrophic lateral sclerosis (**Extended Data Fig. 8b-d**). REEP1 was also found on fragmented LC3-positive mitochondria (**Extended Data Fig. 8e**) under Parkin-mediated mitophagy conditions^33^. Additionally, endogenous REEP2 localized to LC3 punctae in starved SKN-SH cells (**Extended Data Fig. 8f**).

**Fig. 5.**
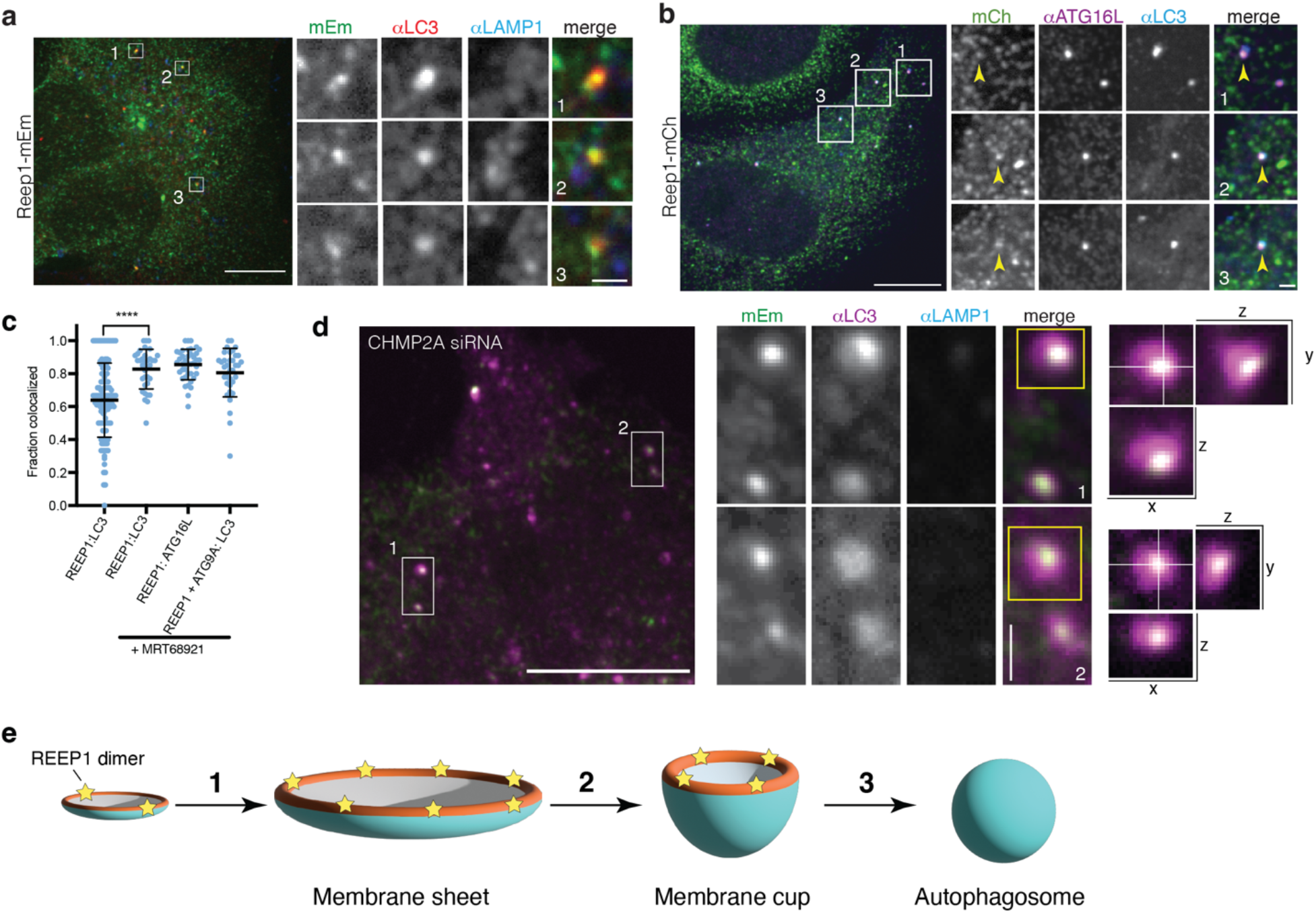
REEP1 proteins accumulate on phagophores in mammalian cells. **a**, U2OS cells stably expressing REEP1-mEm and starved in EBSS for 2h were analyzed by indirect immunofluorescence with antibodies against LC3 (αLC3) and LAMP1 (αLAMP1). Right panels show magnifications of the boxed regions. **b**, As in a, but with cells stably expressing REEP1-mCherry (REEP1-mCh) and immunostained for endogenous ATG16L (αATG16L), an early autophagy marker, and LC3 (αLC3). Arrowheads indicate punctae containing REEP1, ATG16L, and LC3. **c**, Quantification of REEP1 colocalization with autophagosomes stained with anti-LC3, anti-ATG16L, or anti-ATG9A and anti-LC3 in U2OS cells starved in EBSS for 2 h, with or without 1 µM MRT68921 treatment. Shown are means ± standard deviation. Asterisks indicate statistical significance using Student’s *t*-test (*p* < 0.001). **d**, U2OS cells stably expressing Reep1-mEm and treated with CHMP2A siRNA were immunostained for endogenous LC3 and LAMP1. Left panel shows Reep1-mEm and LC3 localization in a 3D projection of a z-series through the cell volume. Middle panels show magnifications of the boxed regions, together with LAMP1 staining. Right panels show orthogonal views of z-series of the indicated LC3 punctae. Scale bars of whole cells, 10 µm; scale bars of magnifications, 1 µm. **e**, Model for the function of REEP1 proteins in autophagosome formation. REEP1 dimers are shown as yellow stars and the phagophore rim is colored in red.

As in *S. pombe*, human REEP1 is recruited early to nascent phagophores where it colocalized with LC3-positive foci that also stained for ATG16L (**Fig. 5b**) and ATG9A (**Extended Data Fig. 9a**), markers of expanding phagophores^34–36^. To further test whether REEP1 joins unclosed phagophores, we treated U2OS cells with the drug MRT68921 to target ULK1 and inhibit phagophore maturation^37^. After starvation, REEP1-mEm colocalized with the majority of LC3 punctae, as well as with ATG16L and ATG9A. The drug also significantly increased colocalization of REEP1 with LC3 (**Fig. 5c**, and **Extended Data Fig. 9b** and **c**). The association of REEP1 with unclosed phagophores is further supported by experiments in which phagophore closure, the terminal step in autophagosome formation^38,39^, was blocked by depletion of the ESCRT-III component CHMP2A (**Extended Data Fig. 9d**). When these cells were starved, LC3 localized to enlarged structures (**Extended Data Fig 9e**) that often contained REEP1-mEm as a smaller focus at the periphery (**Fig. 5d**). This asymmetrical distribution is consistent with REEP1 localizing to the rim of the bending phagophore. Thus, mammalian REEP1 proteins associate with unclosed phagophores, as Rop1 does in *S. pombe*.

## Discussion

Here, we demonstrate that REEP1 family members play an important role in autophagy. Our experiments in *S. pombe* show that the REEP1 homolog, Rop1, is crucial for all forms of autophagy and required for autophagosome formation. It localizes to the rim of expanding phagophores due to its ability to generate high membrane curvature. Experiments with mammalian cells indicate that the function of REEP1 proteins in autophagy is conserved. Based on these results and existing data in the literature, we propose that the REEP1 proteins localize to the rim of the phagophore sheet to stabilize its high membrane curvature (**Fig. 5e**). They would thus be required to keep the two membranes in close proximity. During phagophore growth, the membrane surface area and the rim length of the membrane sheet must initially increase until the size of the double-membrane structure is sufficiently large to enwrap the cargo (**Fig. 5e, step 1**). Thus, the number of REEP1 molecules would increase to stabilize the growing rim. However, at some point the surface area reaches a plateau and the membrane sheet needs to bend to form the quintessential cup-structure around the cargo. Bending implies that the length of the sheet rim, and thus number of REEP1 molecules, must now decrease (**Fig. 5e, step 2**). Eventually, the phagophore double membrane closes on itself by ESCRT-III-mediated fission to generate an autophagosome (**Fig. 5e, step 3**).

How the REEP1 molecules would be removed from the sheet rim is unclear, but perhaps they bud off in vesicles. Mammalian REEP1 proteins, as well as a portion of *S. pombe* Rop1, indeed localize to small vesicles. However, the composition of these vesicles remains to be identified, and it is possible that the vesicles derive from the ER, as the REEP1 proteins partially colocalize with ER markers in *S. pombe* and in mammalian cells. Our model predicts that autophagosomes with closely apposed membranes cannot form in the absence of REEP1 proteins, but some cargo might still occasionally be encapsulated in aberrant autophagosomes, explaining why the absence of Rop1 does not entirely block some forms of autophagy. Our results show that REEP1 is not a functional homolog of Atg40 in *S. cerevisiae*, which has a more limited role as an ERphagy receptor. How phagophores form in budding yeasts, where REEP1 homologs are absent, remains to be investigated.

Finally, our results raise the possibility that autophagy defects may underly the disease pathologies of HSP and the related disorder HMN5B caused by mutations in REEP1 or REEP2^27,28,40^. Several REEP1 disease mutations map to its curvature-generating APH and thus may compromise autophagy.

## Acknowledgments

We thank J. Waters and the Nikon Imaging Center, and M. Ericsson and the Electron Microscopy Facility, both at Harvard Medical School, for technical assistance and use of microscopes. We thank D. Moazed, J.Q. Wu, S. Oliferenko, M. Davidson, and N. Mizushima for strains and plasmids, and D. Pellman, W. Harper, D. Zhao, M. Bao, M. Skowyra, and Z. Ji for critical reading of the manuscript.

## Authors contributions

N.W. initiated the work on REEPs in autophagy. Y.S. identified Rop1 in fission yeast and performed initial experiments with *S. pombe*. N.W. performed all subsequent experiments with yeast Rop1 and the biochemical work. Y.S. performed all experiments with mammalian cells. J.A.P. and S.P.G. performed quantitative mass spectrometry. T.A.R. supervised the project, and Y.S., N.W. and T.A.R. wrote the manuscript.

## Methods

### Fission yeast strain construction and autophagy analysis

**Supplementary Table 1** lists *S. pombe* strains used in this study. Standard genetic and PCR-based gene targeting methods, using pFA6a integration and tagging cassettes, were used to transform yeast cells and to construct strains^1^. All knock-out strains have the coding sequence completely removed from their genomic loci. C-terminally tagged Yop1, Rtn1, Tom20, Pex11, Sfb3, Sec24, Tdh1, Hsc1, Pyk1, Pgk1, and Rop1 were expressed under the control of their endogenous promoters from their chromosomal loci. N-terminally tagged Atg8 was expressed either under the control of the inducible nmt1 or 41nmt1 promoters or expressed from its native promoter, all from its native chromosomal locus. To perform time-lapse imaging of Rop1-mNG and mCherry-Atg8, C-terminally tagged Rop1 was expressed from the inducible 81nmt1 promoter to achieve moderate overexpression. To express YFP, EYFP under the control of the nmt1 promoter was inserted into pJK148 (**Supplementary Table 2**) and integrated into the *leu1* locus. Double or triple mutant strains were generated by genetic crossing and random sporulation. Double and triple mutants containing *fsc1Δ* were constructed by two rounds of transformation because of lower mating efficiency. *S. japonicus* strains were generated as previously described^2^.

To test for autophagy defects by microscopy analysis or proteolytic cleavage assays, *S. pombe* colonies from a freshly streaked plates was grown overnight in liquid Edinburgh minimal medium (EMM5S with 2% glucose, Sunrise Science) at 30°C. The next day (Day 1), cultures were diluted to 0.2 OD600 early AM and late PM to establish cells in logarithmic growth phase. Cultures were subsequently diluted to OD600 of 0.5 on Day 2 and used for analysis. For stationary phase analysis, cells were grown without dilution for another 32-36 h at 30 °C. For DTT treatment, a final concentration of 10 mM DTT was added to the diluted cells and the cells were grown at 30°C for the indicated time period. For nitrogen-starvation, cells corresponding to ∼3 OD600 units were washed once with EMM-N medium with 2% glucose (Sunrise Science), resuspended in EMM-N medium at OD600 of 0.5, and grown at 30 °C for the indicated time period. YE5S (yeast extract with five supplements) was used for *S. japonicus* cell culture. Cells were grown in YE5S for ∼24 h at 30 °C with proper dilutions to keep the culture in log phase. 200 ng/µl of rapamycin was added and cells were cultured for another 20 h before imaging.

For imaging, cells corresponding to 1-2 OD600 units were sedimented at 4,000 g for 30 s and most of the supernatant was removed to leave ∼100 µl. The cells were then resuspended for fluorescence imaging. Imaging was carried out with a spinning-disk mounted on a Ti-motorized inverted confocal microscope (Nikon), as previously described^2^. Images were acquired with a cooled CCD camera (Hamamatsu Photonics) controlled with MetaMorph or NIS-Elements. All display and analysis functions were performed on ImageJ. To quantify colocalization of Rop1 and Atg proteins, a 5×5 pixel area (∼0.32 × 0.32 µm) centered at the brightest Atg pixel was drawn on both channels; the complete or partial overlay of the two squares was binned as colocalization.

To assay for fluorescent protein cleavage, cells corresponding to 3-6 OD600 units were pelleted and washed once with cold water containing 1 mM PMSF. The cells were resuspended in 500 µl of cold water and left on ice for 5 min. Then, 75 µl of 1.85M NaOH, 7.5% (v/v) 2-mercaptoethanol were added to lyse the cells. Next, 75 µl of 55% trichloroacetic acid was added to precipitate the proteins. After centrifugation at 16,000 g for 10 min at 4°C, the supernatant was removed, and the pellet was dried at room temperature for 15 min. The pellet was then suspended in 75 µl of sample buffer containing 8 M urea, 200 mM Tris-HCl pH 6.8, 1 mM EDTA, 5% (w/v) SDS, 0.1% (w/v) bromophenol blue, 1.5% (w/v) DTT. The samples were subjected to 4-20% SDS-PAGE and analyzed by immunoblotting with the following primary antibodies: anti-GFP mouse monoclonal antibody (JL-8, Takara), GFP rabbit polyclonal antibody (PABG1, ChromoTek) for mEGFP detection, anti-mCherry mouse monoclonal antibody (1C51, Abcam), and anti-GAPDH mouse monoclonal antibody (GA1R, Abcam). Staining for total protein was carried out with Revert™ 700 total protein stain kit (LICOR).

DTT-induced ER-stress sensitivity assays were performed as previously described^3^. Cells were grown in liquid EMM5S + 2% glucose at 30°C with 10 mM DTT for 3 days. Cells were washed once with fresh EMM5S before plating. Five-fold dilutions of the indicated strains were spotted onto YE5S plates. Plates were incubated at 30°C and imaged after colony formation.

### Quantitative isobaric tag-based proteomics

*S. pombe* cells were nitrogen-starved for 16 h and ∼ 15 OD600 units of cells were pelleted and frozen in liquid N_2_. Three biological replicates for each strain were collected on separate days, and all replicates were processed on the same day. Frozen cell pellets were washed with water and resuspended in 1 ml of lysis buffer containing 8 M urea, 200 mM EPPS pH 8.5 and protease inhibitors (Pierce A32953). Cells were then lysed by bead-beating and the lysates were clarified by centrifugation at 2,000 g for 30 s. The total protein concentration was determined using the Pierce BCA Protein (23227) assay. The sample was reduced with 5 mM Tris(2-carboxyethyl) phosphine (TCEP) for 30 min, alkylated with 10 mM iodoacetamide for 30 min, and quenched with 10 mM DTT for 15 min. Approximately 100 µg of protein were transferred to a new tube for methanol-chloroform precipitation. The pellet was resuspended in 200 mM EPPS, pH 8.5 and digested at room temperature for 14 h with LysC protease at a 100:1 protein-to-protease ratio. Trypsin was then added at a 100:1 protein-to-protease ratio and the reaction was incubated for 6 h at 37°C. Streamlined–TMT labeling and LS-MS were all done following the protocol described in ^4,5^.

The signal-to-noise measurements of peptides assigned to each protein were summed and these values were normalized so that the sum of the signal for all proteins in each channel was equivalent. Each protein abundance measurement was scaled, such that the summed signal-to-noise for that protein across all channels equals 100, thereby generating a relative abundance (RA) measurement (**Supplementary Table 3**). In all experiments, the total number of protein species detected are the same (> 4000). Data analysis was done by Graphpad Prism and the volcano plots were generated at https://huygens.science.uva.nl/VolcaNoseR/.

### Thin-section transmission electron microscopy (TEM)

EM of fission yeast was performed as previously described with some modifications^6^. Cells corresponding to approximately 10 OD600 units were mixed with 1/25 volume of 25% glutaraldehyde and immediately sedimented at 3,000 rpm for 5 min. The cells were washed once with water and resuspended in freshly prepared 4% KMnO_4_. After incubation at 4 °C for 3 h, the cells were pelleted, washed extensively with water, and resuspended in 2% uranyl acetate and left rotating at 4 °C for 16 h. The cells were again washed extensively with water and dehydrated through a graded ethanol series and embedded in Spurr’s resin. Thin sections were examined using the Tecnai G^2^ Spirit BioTWIN operated at an acceleration voltage of 80 kV or JEOL 1200EX. Images were processed with ImageJ. For quantification of filled vacuole (%), vacuole #, and autophagic structure #/cell, cells with similar length and a clear nucleus visualized in the center were chosen to minimize natural variations in different cross-sections of the cell. Circularity of individual autophagic structures was measured by ImageJ. Data were plotted using Graphpad Prism.

Sample preparation for DAB staining EM was performed as in ^7^ with some modifications. ∼15 OD of starved *S. pombe* cells were harvested and washed once with cold water. The cell pellets were spheroplasted by resuspending in 1ml of 1.2M sorbitol containing 10mg/ml of Zymolyase-20T and rotated at room temperature for 10 min. Cells (1ml) were then layered on top of a 200 µl cushion of 4% PFA /0.2% Glutaraldehyde (in 0.1M Sodium Phosphate buffer, pH 7.4) and immediately pelleted. Next, the supernatant was removed and fixed in 4% PFA/0.2% glutaraldehyde for 1 h and then washed with PBS. Half of the fixed spheroplast was stained for 30 min in a 0.05% DAB-HCl solution with 1:1,000 of 30% H_2_O_2_ to obtain an electron-dense staining of the APEX2 tag; the other half was used as a non-treated control. Staining was washed in PBS, then incubated in filtered 6% potassium permanganate for 1 h. Pellets were washed in PBS until the supernatant is clear.

Samples were dehydrated using solutions of increasing ethanol (EtOH) concentration for 15 min each (7%, 30%, 50%, and 70% [once each] and 100% EtOH [twice]). Low speed centrifuge can be applied if necessary. After the 50% EtOH step, samples were additionally incubated in 1% OsO_4_ /50% EtOH for 1 h. Subsequent resin embedding was done using low-viscosity Spurr’s solution.

### Protein purification and *in vitro* reconstitution

*S. japonicus* Yop1, Yop1(Δ142-165), Rop1, Rop1(Δ107-120), Rop1(Δ126-170), human REEP1, REEP1(Δ102-139), and REEP1(L107P) were expressed as SBP fusion proteins in *E. coli* BL21-CodonPlus (DE3)-RIPL cells (Agilent) and purified as previously described for Yop1/REEP5 proteins^2^. All generated plasmids used in this study are listed in **Supplementary Table 2**. *S. japonicus* Rop1 (sjRop1) lipoprotein particles (LPPs) were purified from the non-sedimentable fraction after 1 h of ultracentrifugation in a Ti-45 rotor (Beckman) at 44,000 rpm at 4 °C, as described^2^. The membrane fractions containing the REEP1 proteins were solubilized in N-dodecyl-β-maltoside (DDM) and the proteins were purified, as described^2^. sjRop1 LPPs and the purified REEP1 proteins from the membrane fractions were further purified by size exclusion chromatography (SEC) on Superose 6 or Superdex 200 (GE Healthcare) columns, respectively.

To incorporate DDM-solubilized proteins into liposomes, protein-free liposomes were first generated with an *S. cerevisiae* Yeast Polar Lipid extract (Avanti) and protein was then added at a 1:300 (protein:lipid) molar ratio with DDM supplemented to a final concentration of ∼0.1%. The mixture was incubated with gentle rotation at 4 °C for 1 h. The detergent was then removed by four successive additions of Bio-Beads SM-2 Resin (Bio-Rad) over the course of ∼24 h at 4 °C. Insoluble aggregates were removed by centrifugation.

### Negative-stain EM

Five µl of reconstituted proteoliposomes or purified sjRop1 LPPs were added to a glow-discharged carbon-coated copper grid (Pelco, Ted Pella Inc.) for 1 min. Excess sample was blotted off with filter paper. The grids were washed once with deionized water, and then stained once with freshly prepared 1% uranyl acetate for 30 s prior to blotting and air-drying. Images were collected using a Tecnai G^2^ Spirit BioTWIN transmission electron microscope operated at an acceleration voltage of 80 kV, or a JEOL 1200EX microscope equipped with an AMT 2k CCD camera.

### Site-specific photo-crosslinking

To incorporate photoreactive Bpa probes into sjRop1, amber codons were introduced into Rop1 at various positions (see **Supplementary Table 2** for plasmids). The mutated Rop1 expression plasmid and the amber suppressor tRNA plasmid were co-transformed into BL21 (DE3) cells (NEB), as described^2^. One mM final concentration of H-Bpa-OH (Chempep) was added to the medium, and the amber suppressor tRNA was induced 1–2 h before induction of sjRop1 expression.

For photo-crosslinking, 20 µl of a membrane suspension in 25 mM HEPES/KOH pH 7.5, 150 mM NaCl was added to Thermowell PCR tubes (Corning) and placed onto an ice-cold metal block. The samples were irradiated for 30 min with a long-wave UV lamp (Blak-Ray) and then analyzed by SDS-PAGE and western blotting with DyLight 800-labeled streptavidin (Invitrogen) diluted to 1:4000 in 1% BSA.

### Purified Rop1 and Atg8 pull-downs

To test for interactions between Rop1 and Atg8 family proteins, *S. cerevisiae* Atg8 was tagged at the N-terminus with His10 and *S. pombe* Atg8 was tagged at the N-terminus with GST (see **Supplementary Table 2**). The proteins were expressed in *E. coli* and purified on Ni-NTA and GSH beads, respectively, essentially as described^8^. The proteins were further purified by SEC on a Superdex 75 (GE Healthcare) column. SBP-tagged full-length sjRop1, scAtg40, or mutants with mutated putative AIM motifs [sjRop1(AIM) and scAtg40(AIM)] were expressed in *E. coli*.

The DDM-solubilized membrane fractions were incubated with streptavidin beads and washed extensively with wash buffer containing 25 mM HEPES/KOH pH 7.4, 150 mM NaCl, 0.05% DDM, and protease inhibitors. The beads were then incubated with purified Atg8 in buffer containing 0.05% DDM for 1 h at 4 °C. The beads were washed again with 20 column volumes (CV) of buffer and eluted with one CV of 2x Laemmli sample buffer (Bio-Rad). Thirty % of the eluted sample was subjected to SDS-PAGE gel and Coomassie blue staining. A sample of the input material (5%) was analyzed in parallel.

### Mammalian cell culture

Human U2OS (original gift from J, Blenis) and SKN-SH cells (ATCC) were maintained at 37 °C and 5% CO_2_ in DMEM or EMEM, respectively, supplemented with 10% FBS and penicillin-streptomycin (100 U/ml). REEP1-mEm, REEP1-mCherry, and RFP-Sec61β stable U2OS lines were generated by transfecting the respective gene-encoding plasmids and selecting with 0.5 µg/ml G418 to establish pooled lines, and clonal lines were then established by single cell isolation. HA-Parkin expressing REEP1-mEm U2OS cells were established by retroviral infection of the REEP1-mEm clonal U2OS line and selected with puromycin (1 µg/ml). pHUJI-LC3B expressing SKN-SH cells were established by lentiviral infection and hygromycin selection (0.2 µg/ml). All cell lines were routinely verified to be mycoplasma-free by DAPI staining.

For transient plasmid transfections, cells were plated into 6-well plates so that they were ∼75% confluent at the time of transfection. 1-2 µg of total plasmid DNA was transfected using lipofectamine 3000 (Thermo Scientific). Cells were trypsinized and replated onto No 1.5 coverglass the following day and analyzed ∼12-16 h later by indirect immunofluorescence. CHMP2A RNAi experiments were performed using Dharmacon SMARTpool siRNAs (human CHMP2A #L-020247-01, or negative control #D-001810-10-05, Horizon), where 20 nM of oligonucleotides were reverse transfected into cells using RNAiMax (Thermo Scientific) and analyzed 48 hours later. For starvation experiments, cells were washed in EBSS (Millipore Sigma #E3024) three times before incubation in this medium for the indicated time. For mitophagy induction, 10 µM of antimycin A (Millipore Sigma #A8674) and 5 µM of oligomycin A (Millipore Sigma #75351) were added in DMEM to HA-Parkin expressing U2OS cells for the indicated time period. For ULK1 inhibition, 1 µM of MRT68921 (Millipore Sigma #SML1644) was used.

### Mammalian DNA plasmids, antibodies, and reagents

Human REEP1-HA, REEP2-HA, REEP3-HA, and REEP5-HA plasmids were gifts from C. Blackstone and have been described in ^9^. REEP1-mEmerald and REEP1-mCherry plasmids were generated by inserting a codon-optimized human REEP1 gblock into mEmerald-N1 (gift from M. Davidson; Addgene #53976) or mCherry-N1 backbones (Clontech). The REEP4-mCherry plasmid was constructed by inserting a codon-optimized human REEP4 gblock into mCherry-N1 using Gibson assembly. The FAM134B-HA plasmid was cloned by inserting a codon-optimized gblock encoding human Fam134B with an HA-tag at its C-terminus by Gibson assembly into pGW1. The pHUJI-LC3B construct was cloned by inserting a codon-optimized gblock encoding pHUJI fused to the N-terminus of human LC3B into the NotI site of pENTR4 (Addgene #17424), and then Gateway-recombined with LR clonase II (Thermo Scientific) into the pLenti CMV Hygro destination vector (gift from Eric Campeau, Addgene #19066). All gblocks were purchased from IDT.

pMXs-IP HA-Parkin (gift form Noboru Mizushima, #38248) was obtained from Addgene. GFP-Sec61β and RFP-Sec61β have been described previously in ^10^; SOD1(A4V)-GFP has been described in ^11^.

The following antibodies were used: Monoclonal rat anti-HA (3F10, Millipore Sigma #11867423001); polyclonal rabbit anti-Reep4 (Millipore Sigma/Atlas #HPA042683); monoclonal mouse anti-Reep2 (clone S326D-2, Thermo Scientific); polyclonal rabbit anti-Rtn4a/b (Abcam #ab47085); monoclonal mouse anti-LAMP1 (H4A3, Millipore Sigma #MABC1108); polyclonal rabbit anti-LC3 (MBL #PM036); monoclonal mouse anti-LC3 (MBL #M152-3); monoclonal rabbit anti-panGABARAP (Abcam #ab109364); monoclonal mouse anti-P62 (BD Biosciences, #610832); polyclonal rabbit anti-ATG9A (MBL #PD042); polyclonal rabbit anti-ATG16L (MBL #PM040); polyclonal rabbit anti-CHMP2A (Thermo Scientific #PA5-100092); and anti-cytochrome c (BD Biosciences #556432).

### Indirect immunofluorescence, microscopy, and immunoblotting of mammalian cells

Indirect immunofluorescence was performed using standard procedures. U2OS cells were grown on acid washed No 1.5 glass coverslips, fixed in PBS/4% paraformaldehyde (Electron Microscopy Services), and permeabilized with PBS/0.1% TritonX-100 for 5 min. Samples were blocked in PBS/1% FBS/0.01% TX-100 for 1 h, and then sequentially probed with primary and secondary antibodies conjugated to Alexafluor dyes (405, 488, 568, or 647 from Thermo Scientific) with 3x PBS washes after each antibody treatment. Samples were mounted onto glass slides with fluoromount-G (Southern Biotech).

Confocal fluorescence imaging was performed using the same Nikon imaging system described for yeast, with Nikon LUN-F XL solid state lasers (405, 488, 561, and 640 wavelengths) and a 100X Plan Apo 1.4 N/A oil objective. FIJI/ImageJ was used for all imaging analysis and display functions. For display, fluorescence intensities of raw 16-bit images were linearly scaled across the entire image, saved as 8 bit, and pseudo-colored. Images showing cropped enlargements were similarly processed. For quantification correlating REEP1 expression levels with localization phenotypes, images were taken with a 63X 1.4 N/A oil objective using identical exposure settings for the anti-HA fluorescence channel, and the average fluorescence intensity, subtracted for background, of each cell was compared to its localization. For quantification of ATG16/LC3 foci colocalization, images were taken at 100X with the same exposure settings and ATG16 or LC3 positive foci were identified by thresholding intensities. Co-presence of a REEP1 punctum (fluorescence signal above background) to these structures was quantified as colocalized. Pearson’s correlation coefficients were calculated on images taken at 100X on an ∼8×8 µm area at the cell periphery where the ER was a clear monolayer using the JaCoP plugin. All quantifications were performed on raw 16-bit images. All calculations, statistics and graphs were analyzed and plotted using Microsoft Excel and Graphpad Prism.

Immunoblotting was performed by lysing cells in modified RIPA buffer (50 mM Hepes pH 7.4, 150 mM NaCl, 1 mM MgCl_2_, 1% TX-100, 0.1% deoxycholate, 0.1% SDS, with protease inhibitors). Clarified lysates were denatured in Laemmli SDS buffer and resolved on a 4-20% gradient TGX PAGE gel (BioRad). Blots were transferred and visualized using HRP-conjugated secondary antibodies and western lightning ECL (GE) on an Amersham 600 Imager (GE) or with Revert 520 total protein stain (Licor) on a LICOR M system.

---

**Extended Data Fig. 1.**
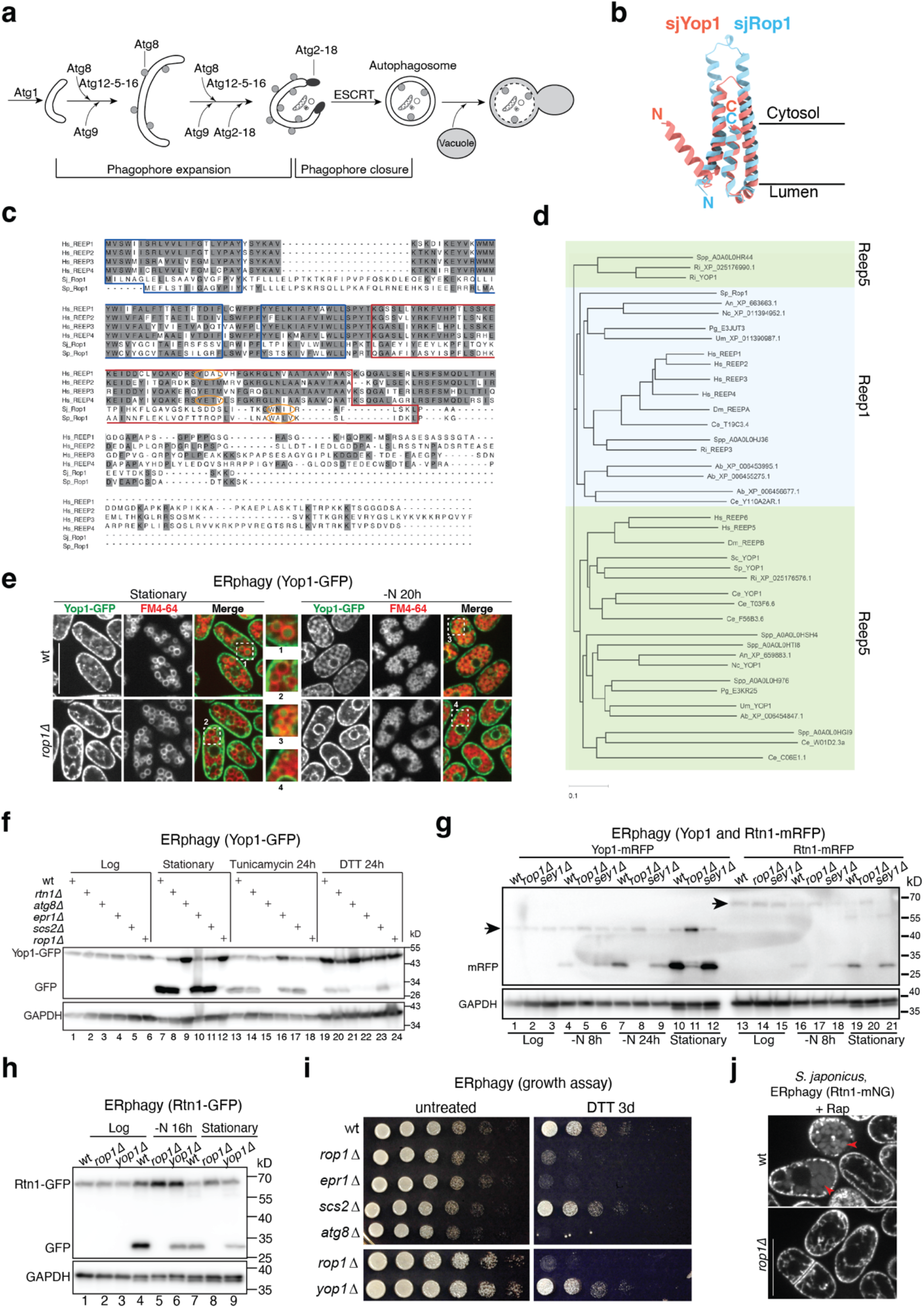
Rop1 is required for ERphagy in fission yeasts *S. pombe* and *S. japonicus*. **a**, Steps of autophagy in yeast. The process is conserved in mammals, with vacuoles replaced by lysosomes, Atg1 by ULK1, and Atg8 by LC3 and GABARAP. **b**, Side views of superimposed cartoon models of *S. japonicus* (sj) Rop1 and Yop1, predicted by Alphafold^1^. The APHs of both proteins were omitted for clarity. **c**, Sequence alignment of human REEP1-4 proteins and *S. japonicus* and *S. pombe* Rop1 using ClustalW. Identical residues are highlighted in gray. Predicted TM, APH, and AIM regions are outlined in blue, red, and orange, respectively. **d**, The phylogeny of REEP proteins was analyzed using ClustalW. Unique gene-encoded REEP protein sequences from representative species of all fungal phyla (Sp, *S. pombe;* Sc, *S. cerevisiae;* Ri, *R. irregularis;* Nc, *N. crassa;* An, *A. nidulans;* Um, *U. maydis;* Ab, *A*.*bisphoru*s; Spp, *S. punctaetus*; Pg, *P. graminis*) were compared with REEP proteins from human (Hs), *D. melanogaster* (Dm), and *C. elegans* (Ce). The REEP5/Yop1 and REEP1 subfamilies are indicated in green and blue, respectively. Note that all species, except *S. cerevisiae*, possess at least one distinct REEP1-like protein. **e**, The ER protein Yop1 was tagged with GFP (Yop1-GFP) and expressed in wild-type (wt) or *rop1Δ S. pombe* cells. Cells in stationary phase or deprived of nitrogen for 20h (-N 20h) were analyzed by fluorescence microscopy. The vacuoles were stained with FM4-64. The right panels show merged images, and boxed regions 1-4 are shown in magnified views. Scale bar, 10 µm. **f**, Yop1-GFP was expressed in wt or mutant cells. Lysates from cells in logarithmic (Log) or stationary phase, as well as from cells treated with tunicamycin or DTT for 24h, were analyzed by SDS-PAGE and immunoblotting with GFP antibodies. Blotting with GAPDH antibodies served as loading control. Note that Scs2 is a component in the Epr1 pathway^2^ and its absence does not affect cleavage of Yop1. **g**, As in f, but with Yop1-mRFP and Rtn1-mRFP. Arrows point to the positions of the full-length proteins. **h**, As in f, but with Rtn1-GFP. **i**, wt or mutant *S. pombe* cells were treated with DTT for 3 days and plated after serial dilution. Controls were performed with untreated cells. **j**, *S. japonicus* cells expressing Rtn1-mNeonGreen (Rtn1-mNG) in wt or *rop1Δ* were treated with 200 ng/µl rapamycin in YES medium for 20 h before imaging. Arrowheads point to Rtn1-mNG in vacuoles. Scale bar, 10 µm.

**Extended Data Fig. 2.**
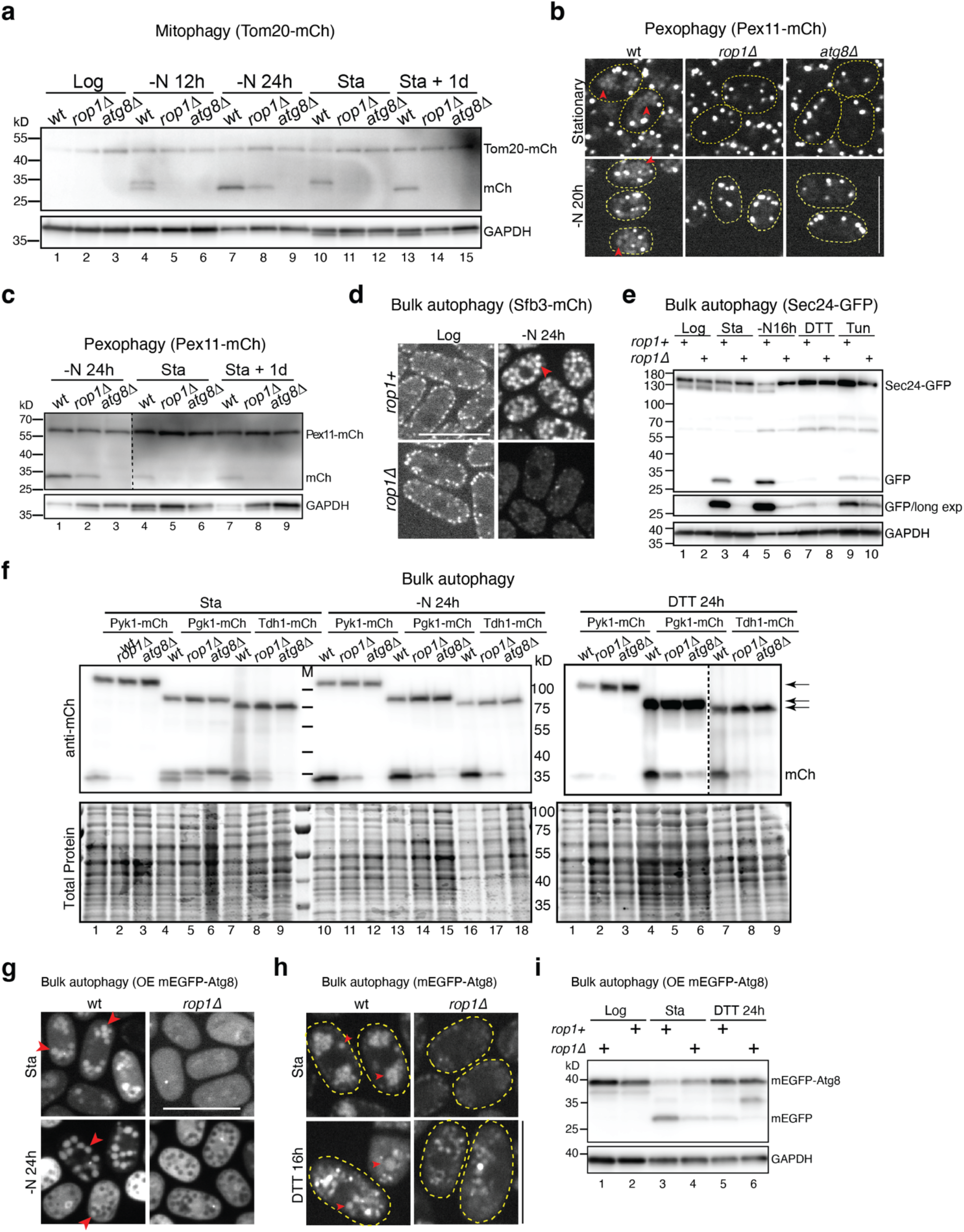
Rop1 is involved in mitophagy, pexophagy, and bulk autophagy in *S. pombe*. **a**, The mitochondrial protein Tom20 was tagged with mCherry (Tom20-mCh) and expressed in wild-type (wt) or mutant cells. The cells were analyzed in logarithmic (Log) or stationary (Sta) phase or after nitrogen starvation for 12h or 24h (-N 12h or -N 24h). Some cells were incubated for an additional day after reaching Sta phase (Sta + 1d). In all cases, cell lysates were analyzed by SDS-PAGE and immunoblotting with mCherry antibodies. Blotting with GAPDH antibodies served as loading control. **b**, The peroxisomal protein Pex11 was tagged with mCherry (Pex11-mCh) and expressed in wt and mutant cells. The cells were nitrogen-starved for 20h (-N 20h) and analyzed by fluorescence microscopy. Red arrowheads point to Pex11-mCh in vacuoles. **c**, As in a, but for Pex11-mCh. **d**, A mCherry fusion of Sfb3 (Sfb3-mCh), a component of the cytosolic COPII complex, was expressed in wt or *rop1Δ* cells. The cells were analyzed in Log phase, or after nitrogen starvation for 24h (-N 24h), by fluorescence microscopy. Red arrowhead points to Sfb3-mCh in vacuoles. **e**, As in **a**, but with the COPII component Sec24 tagged with GFP (Sec24-GFP). A long exposure (exp) of the immunoblot is shown as well. **f**, As in a, but for mCherry fusions of the cytosolic proteins Pyk1 (Pyk1-mCh), Pgk1 (Pgk1-mCh), and Tdh1 (Tdh1-mCh). Arrows point to the full-length proteins. As a loading control, the blot was stained for total protein with Revert700 (lower panels). **g**, As in b, but for the overexpressed autophagy component Atg8 tagged with mEGFP under the inducible *nmt1* promotor (OE mEGFP-Atg8). **h**, As in g, except that mEGFP-Atg8 was expressed under its native promotor. The boundaries of the cells are indicated by dotted lines. **i**, As in a, but for overexpressed mEGFP-Atg8. Scale bars, 10 µm.

**Extended Data Fig. 3.**
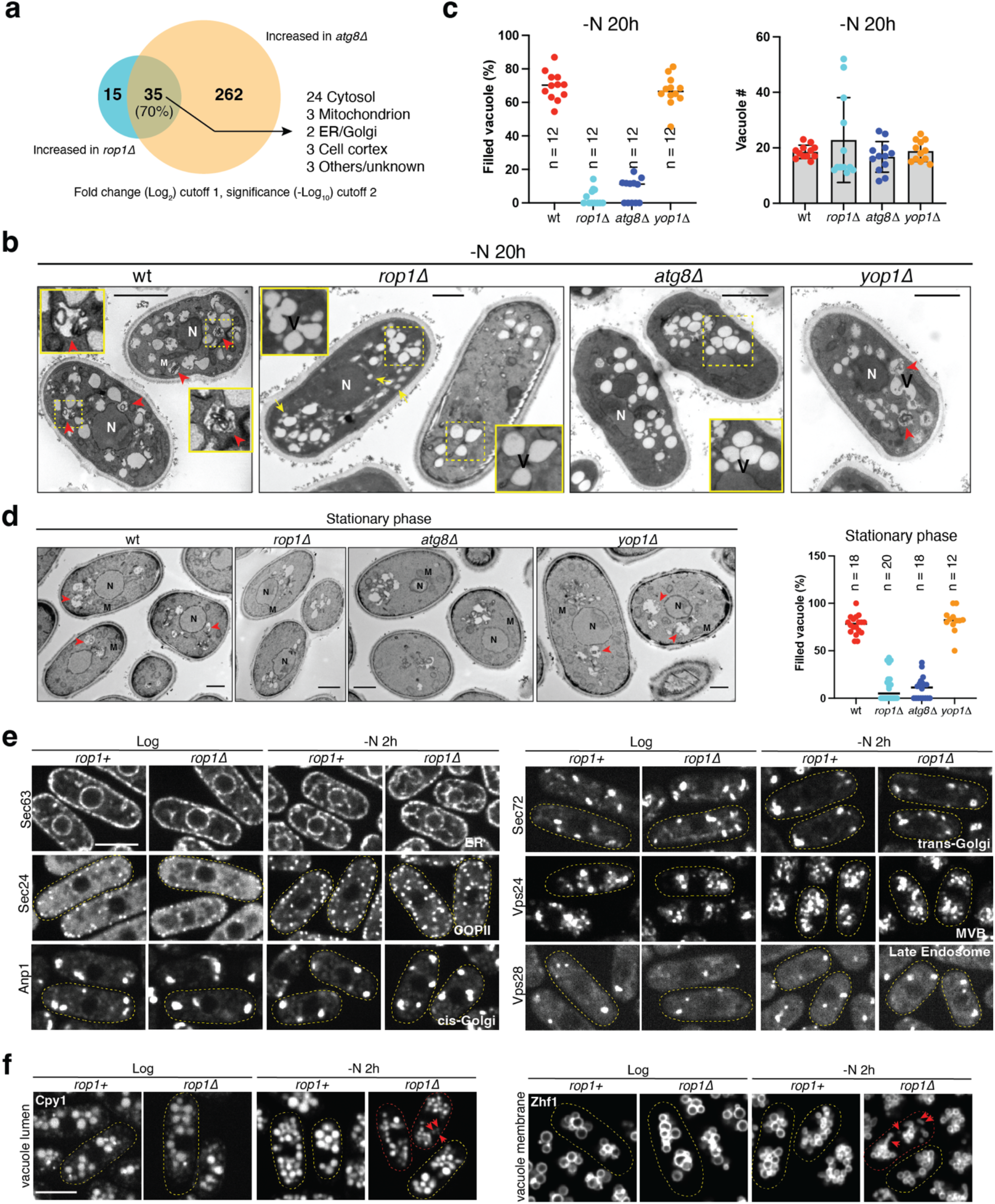
Rop1 is involved in autophagosome formation. **a**, Protein accumulation in nitrogen-starved Rop1- or Atg8-lacking cells was determined by quantitative isobaric tag-based proteomics. The Venn diagrams show the accumulating proteins and their overlap. Gene Ontology (GO) annotations of the overlapping proteins are listed on the right. **b**, TEM was performed with nitrogen-starved (-N 20h) wild-type (wt) and mutant cells. Red arrows point to vacuoles filled with electron-dense autophagic cargo, likely residual membranes of autophagosomes. Yellow arrows point to unusually small vacuoles seen in *rop1Δ* cells. N, nucleus; V, vacuole. Scale bar, 1 µm. **c**, Quantification of the results in b. Shown is the percentage of filled vacuoles per cell (left panel; the lines indicate the means) and the total number of vacuoles per cell (right panel; means and standard deviations). n, number of cells analyzed. **d**, As in b, but for cells in stationary phase. M, mitochondrion. Shown on the right is percentage of filled vacuoles (lines indicate the means). **e**, Markers of the bulk ER (Sec63), ER exit sites (Sec24), cis-Golgi (Anp1), trans-Golgi (Sec72), multivesicular bodies (Vps24), or late endosomes (Vps28) were tagged with fluorescent proteins and expressed in *rop1*^*+*^ and *rop1Δ* cells. The cells were analyzed in Log phase or after starvation for 2h by confocal fluorescence microscopy. The dotted line shows the boundaries of cells. Scale bar, 5 µm. **f**, As in e, but with markers of the vacuole lumen (Cpy1) or membrane (Zhf1). Red arrowheads point to abnormally small vacuoles.

**Extended Data Fig. 4.**
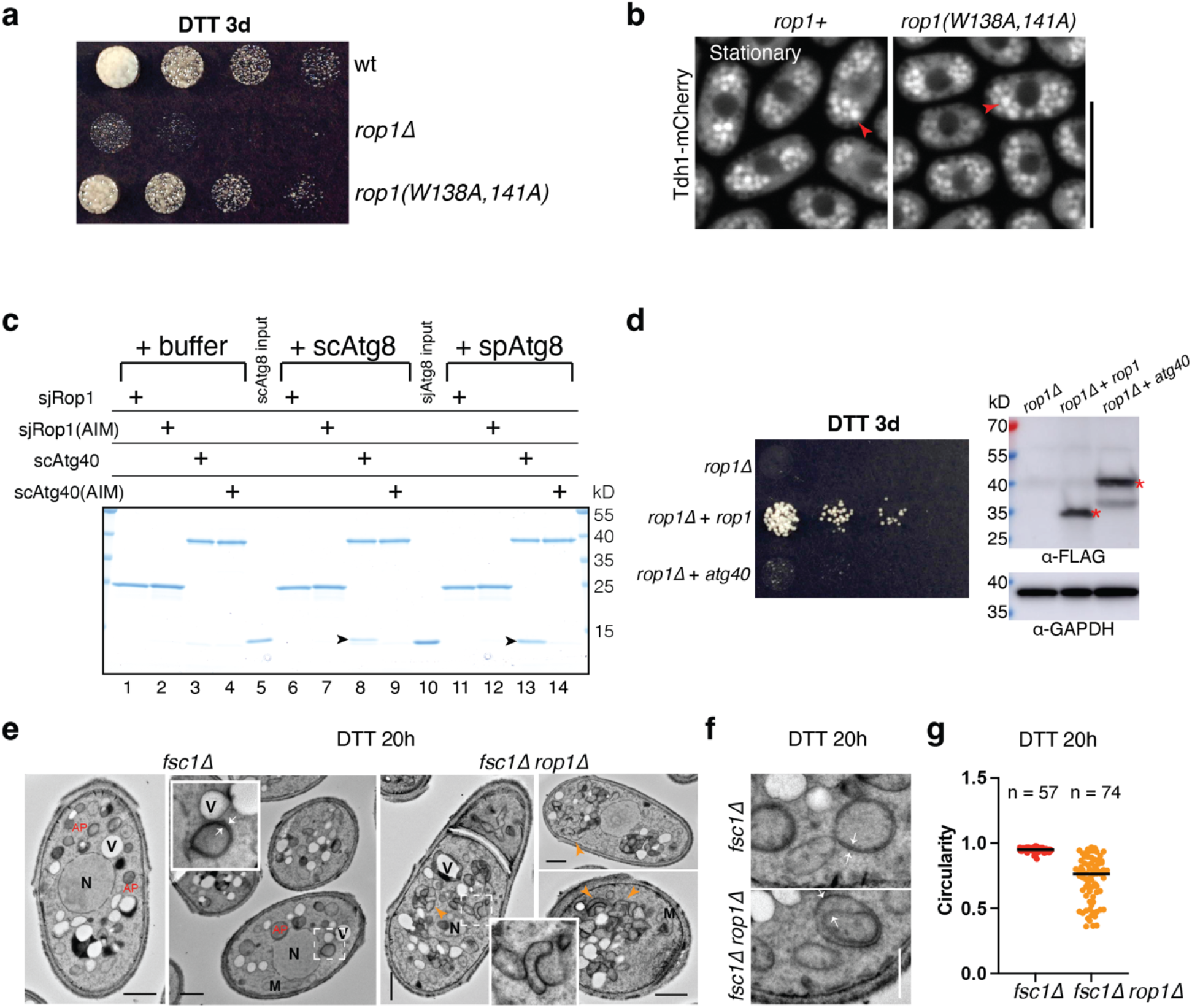
Rop1 is not an ERphagy receptor. **a**, Wild-type (wt) cells or the indicated mutant cells were treated with DTT for 3 days and plated after serial dilution. The Rop1 (W138, 141A) mutant has an abrogated AIM motif. **b**, The cytosolic protein Tdh1 was tagged with mCherry (Tdh1-mCh) at its genomic locus and expressed in wt or rop1(W138, 141A) mutant cells. Stationary phase cells were analyzed by fluorescence microscopy. Red arrowheads point to vacuoles. Scale bar, 10 µm. **c**, *S. japonicus* Rop1 (sjRop1) or *S. cerevisiae* Atg40 (scAtg40), or mutants of these proteins with abrogated AIM motifs [sjRop1(AIM) and scAtg40(AIM)], were tagged with SBP at the C-terminus and purified from *E. coli*. The proteins were incubated with buffer or purified Atg8 from *S. cerevisiae* (scAtg8) or *S. pombe* (spAtg8). The samples were then incubated with streptavidin beads and the bound material analyzed by SDS-PAGE and Coomassie-blue staining. Arrowheads point to co-precipitated Atg8. Input corresponds to 5% used in the pull-down experiments. **d**, *S. cerevisiae* Atg40, tagged with 5 FLAG epitopes (scAtg40-5FLAG), was expressed in *rop1Δ* cells from the endogenous *rop1* locus under the *rop1* promoter (*prop1*). Controls were performed with cells expressing spRop1-5FLAG. The cells were treated with DTT for three days (DTT 3d) and plated after serial dilution (left panel). Cell lysates were also analyzed by SDS-PAGE and immunoblotting with FLAG antibodies (right panel). **e**, *fsc1Δ* or *fsc1Δ rop1Δ* cells were treated with DTT for 20h and analyzed by TEM. AP, autophagosome; N, nucleus; M, mitochondrion; V, vacuole. Scale bar, 1 µm. Orange arrow heads point to irregular autophagic structures. Squares outlined with dashed lines are magnified in the insets. White arrows point to autophagosomes with distinct lipid bilayers. **f**, As in e, showing additional examples of magnified views of autophagic structures. Scale bar, 500 nm. **g**, Quantification of the circularity of autophagic structures shown in e.

**Extended Data Fig. 5.**
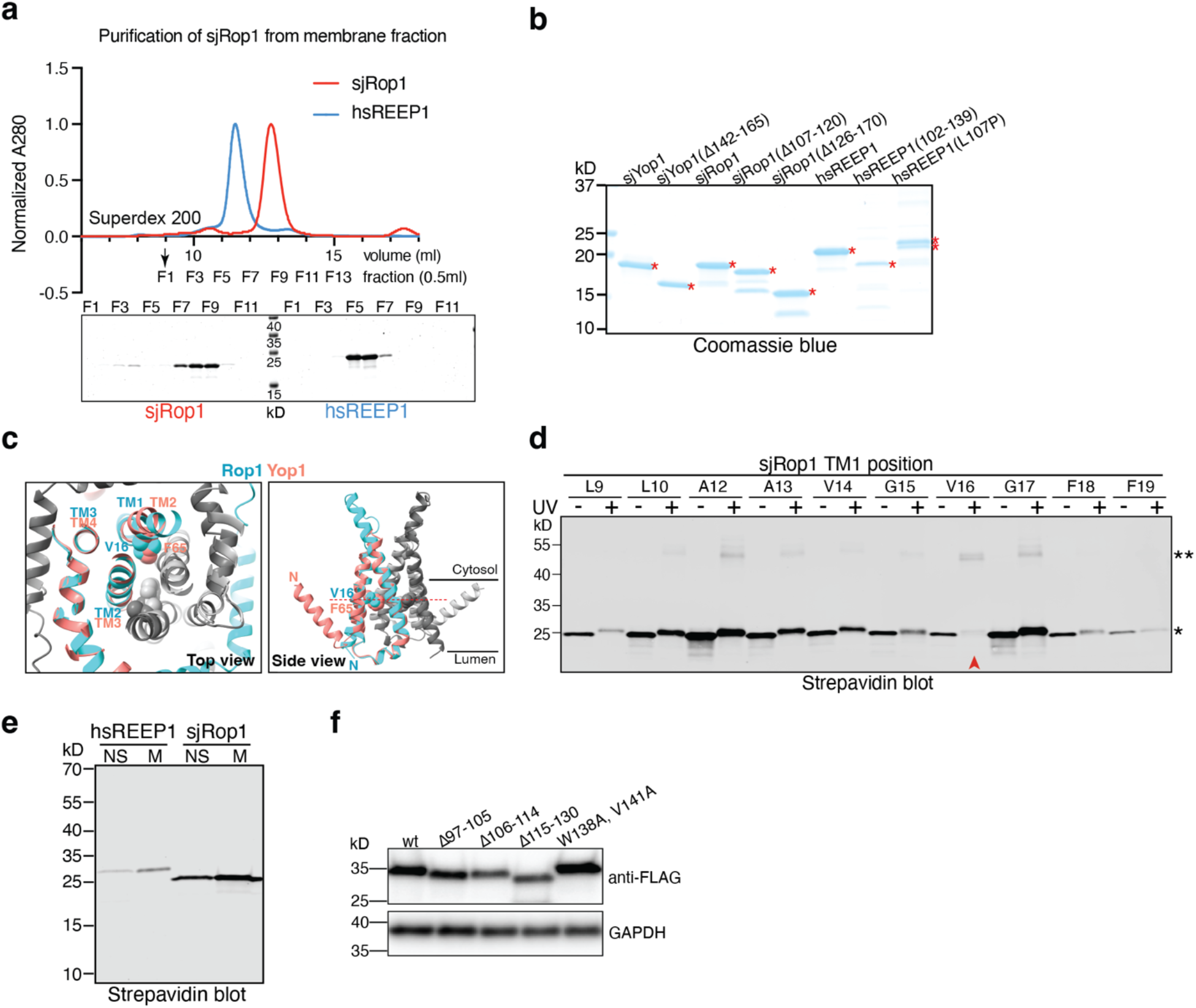
Purification and dimerization of REEP1 proteins. **a**, SBP-tagged sjRop1 or human REEP1 (hsREEP1) were expressed in *E. coli*. A membrane fraction was obtained by ultracentrifugation and solubilized in DDM. The proteins were purified with streptavidin beads and subjected to gel filtration on a Superdex 200 column. The UV absorbance at 280 nm was monitored (upper panel), and fractions were analyzed by SDS-PAGE and Coomassie-blue staining (lower panels). **b**, Wild-type sjRop1 or hsREEP1, or the indicated mutants, were purified and analyzed by SDS-PAGE and coomassie blue staining. Red stars indicate the positions of the purified proteins. Additional bands are likely caused by proteolysis. **c**, Predicted interaction of the monomers of sjRop1 and sjYop1 in the dimers. Shown is the superposition of the TM segments of the two structures in top (cytosolic) and side views, with one monomer in color (Rop1 in blue and Yop1 in pink) and the other in grey. Amino acids V16 of sjRop1 and F65 of Yop1 give the strongest dimer crosslinks (see panel **d**) and are shown as balls. **d**, SBP-tagged sjRop1 was expressed in *E. coli* with photoreactive Bpa probes incorporated at the indicated positions of TM1 by amber codon suppression. Where indicated, a membrane fraction was irradiated with UV light, and the samples were analyzed by SDS-PAGE, followed by blotting with dye-labeled streptavidin and fluorescence scanning. The red arrowhead indicates the position with strongest dimer crosslinks. **e**, SBP-tagged hsREEP1 and sjRop1 were expressed in *E. coli*. Cell lysates were subjected to ultracentrifugation and the membrane (M) and non-sedimentable (NS) fractions analyzed by SDS-PAGE, followed by blotting with dye-labeled streptavidin and fluorescence scanning. **f**, The expression levels of FLAG-tagged wt Rop1 or the indicated mutants was determined in logarithmically growing *rop1Δ* cells by immunoblotting with FLAG antibodies. Blotting with GAPDH antibodies served as a loading control.

**Extended Data Fig. 6.**
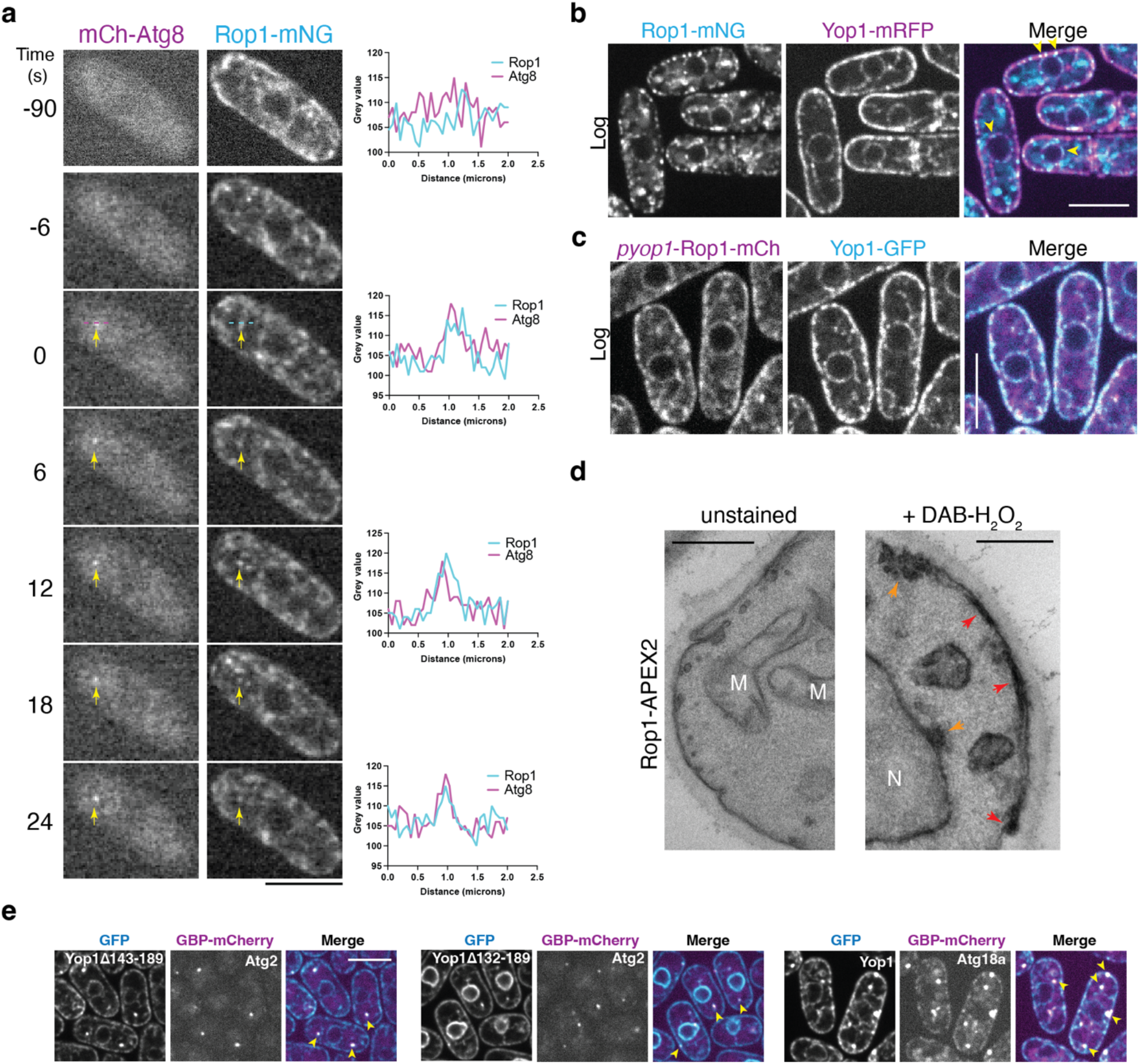
Localization of Rop1 in *S. pombe*. **a**, Rop1 tagged with mNeonGreen (Rop1-mNG) was co-expressed with mCherry-tagged Atg8 (mCh-Atg8) in *S. pombe* cells. The cells were nitrogen-starved for 3h (-N 3h) and analyzed by confocal fluorescence microscopy. Shown is the montage of a time-lapse movie. At time point zero, a mCh-Atg8 punctum (phagophore) appears (arrow). The right panels show line scans across the Atg8 punctum. Scale bar, 5 µm. **b**, Cells expressing Rop1-mNG and Yop1-mRFP at endogenous levels from their genomic loci were visualized in logarithmic (Log) phase. Arrowheads point to colocalization of the two proteins in the ER. Scale bar, 5 µm. **c**, As in b, but Rop1-mCherry was overexpressed under the *yop1* promoter from the *leu1* genomic locus (*pyop1*-Rop1-mCh) while Yop1 was tagged with GFP and expressed from its genomic locus (Yop1-GFP). **d**, Cells expressing Rop1-APEX2 were nitrogen-starved for 2h, fixed, and stained without (left) or with (right) DAB and H_2_O_2_ for 20 min before preparation for TEM. M, mitochondria; N, nucleus. Red arrowheads point to staining of the peripheral ER and orange arrowheads to staining of vesicles near the ER and nuclear envelope. Scale bar, 500 nm. **e**, Atg2 or Atg18a was tagged with GBP-mCherry and coexpressed with mutant or wild-type Yop1-GFP (see also Fig. 4k). The cells were imaged after 2h of nitrogen starvation. The arrowheads point to colocalization between the Atg proteins and Yop1. The Yop1Δ143-189 and Yop1Δ132-189 mutants contain and lack the APH, respectively. Scale bar, 5 µm.

**Extended Data Fig. 7.**
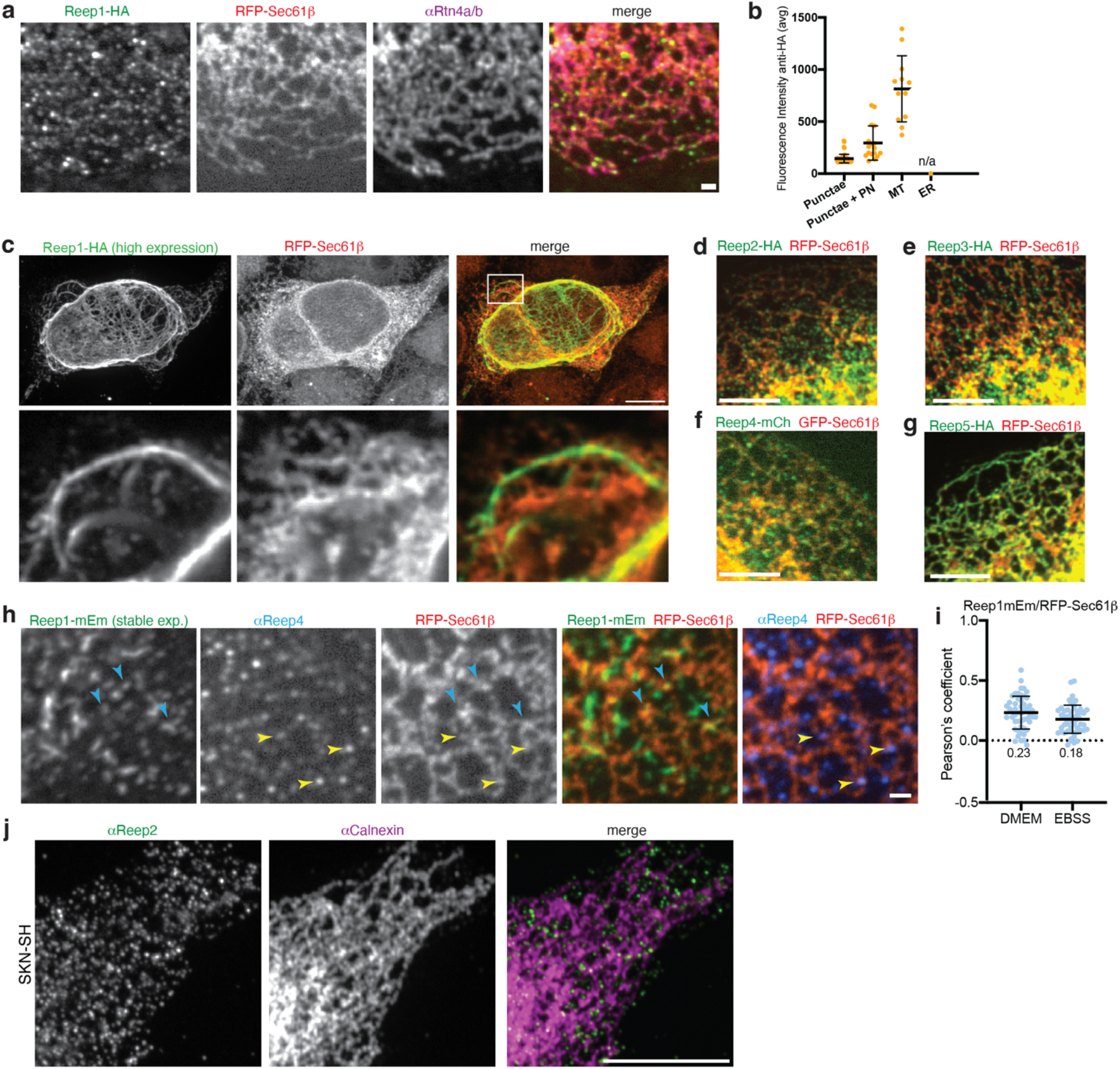
Localization of human REEP1 proteins. **a**, Visualization of transiently transfected HA-tagged REEP1 (REEP1-HA) in U2OS cells. The localization of REEP1-HA (αHA) is compared with that of stably expressed RFP-Sec61β, a general ER marker, and endogenous Rtn4 (αRtn4a/b), a tubular ER marker. Note the vesiculated appearance of REEP1 and limited overlap with bulk ER. Similar results were obtained for the other human REEP1 subfamily members REEP2, REEP3, and REEP4 (see d-j). **b**, Quantification of different localization patterns of transiently transfected REEP1-HA cells. The previously observed elongated membrane tubules described as microtubule-bundled ER^3^ were only generated in highly expressing, transiently transfected cells (see also c). PN, bright perinuclear localization; MT, bundled microtubule phenotype. **c**, As in a, but in a cell with high expression of REEP1-HA showing the microtubule-bundled phenotype quantified in b. Top row shows a maximal projection of a z-stack. Bottom row shows a magnified view of a single confocal slice of the boxed region. Note that REEP1-HA poorly colocalizes with the bulk ER. **d**, As in a, but with cells transfected with REEP2-HA. **e**, As in a, but with cells transfected with REEP3-HA. **f**, As in a, but with stably expressing GFP-Sec61β cells transfected with REEP4-mCherry. **g**, As in a, but with cells transfected with REEP5-HA, an ER shaping protein. Note the punctate appearance of REEP2-4 compared to the tubular ER localization of REEP5. **h**, As in a, but with stably expressing REEP1-mEmerald (REEP1-mEm) cells transiently transfected with RFP-Sec61β, and co-stained for endogenous REEP4 (αREEP4). Arrowheads indicate a few REEP1 (blue arrows) and REEP4 (yellow arrows) punctae that colocalize with ER. **i**, Pearson’s correlation coefficients between REEP1-mEm and RFP-Sec61β in cells as in h under fed (DMEM) or starved (EBSS) conditions. Shown are mean ± standard deviations (*n* = ∼50 cells). Note that REEP1 shows limited colocalization with the bulk ER. **j**, Visualization of endogenous REEP2 (αReep2) and of the general ER marker calnexin (αCNX) in SKN-SH cells, which express high levels of endogenous REEP2 (https://www.proteinatlas.org/search/REEP2)^4^. Scale bars, a, h, 1 µm; c, j, 10 µm; d-g, 5 µm.

**Extended Data Fig. 8.**
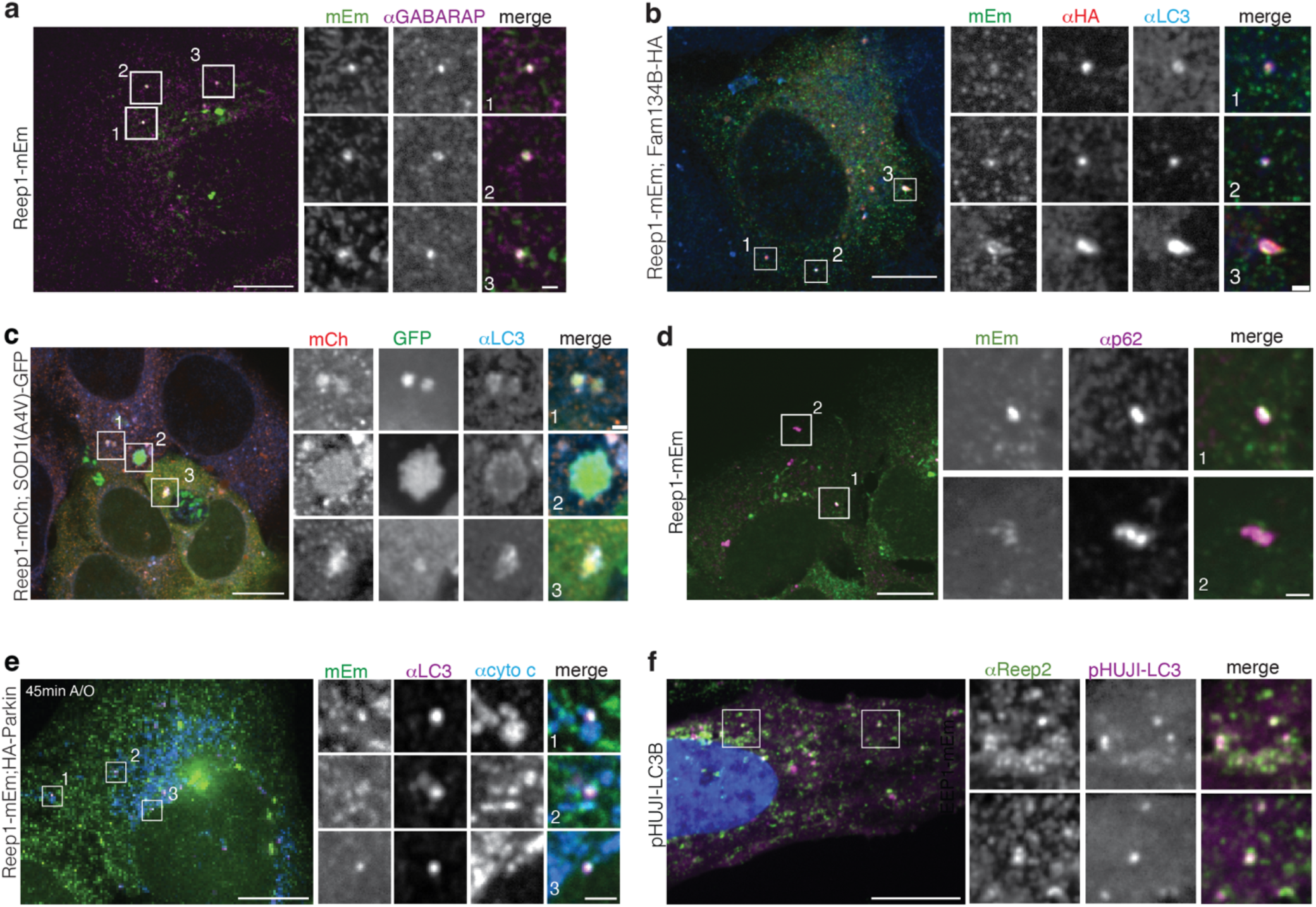
REEP1 proteins colocalize with autophagic receptors and cargo. **a**, U2OS cells stably expressing REEP1-mEm and starved in EBSS for 2h were analyzed by indirect immunofluorescence with a pan-GABARAP antibody. Right panels show magnifications of the boxed regions. **b**, As in a, but with REEP1-mEm expressing cells transfected with HA-tagged FAM134B, an ERphagy receptor, and immunostained with anti-HA and anti-LC3 antibodies. **c**, U2OS cells stably expressing REEP1-mCherry (REEP1-mCh) and transiently transfected with SOD1(A4V)-GFP were immunostained with anti-LC3 antibodies. Shown are cells with SOD1 aggregates. **d**, As in a, but with cells starved for 6 h in EBSS and immunostained for endogenous p62 (αp62), an autophagic cargo receptor. **e**, U2OS cells stably expressing REEP1-mEm and HA-tagged Parkin were treated with antimycin and oligomycin for 45 min to induce selective mitophagy and immunostained with anti-LC3 and the mitochondrial marker anti-cytochrome c (αcyto c). **f**, Colocalization of endogenous Reep2 (αReep2) with stably expressed LC3 fused to the pH-sensitive fluorescent protein pHUJI (pHUJI-LC3B) in SKN-SH cells starved in EBSS for 1 h. Scale bars for all left panels showing whole cells, 10 µm; for all magnifications, 1 µm.

**Extended Data Fig. 9.**
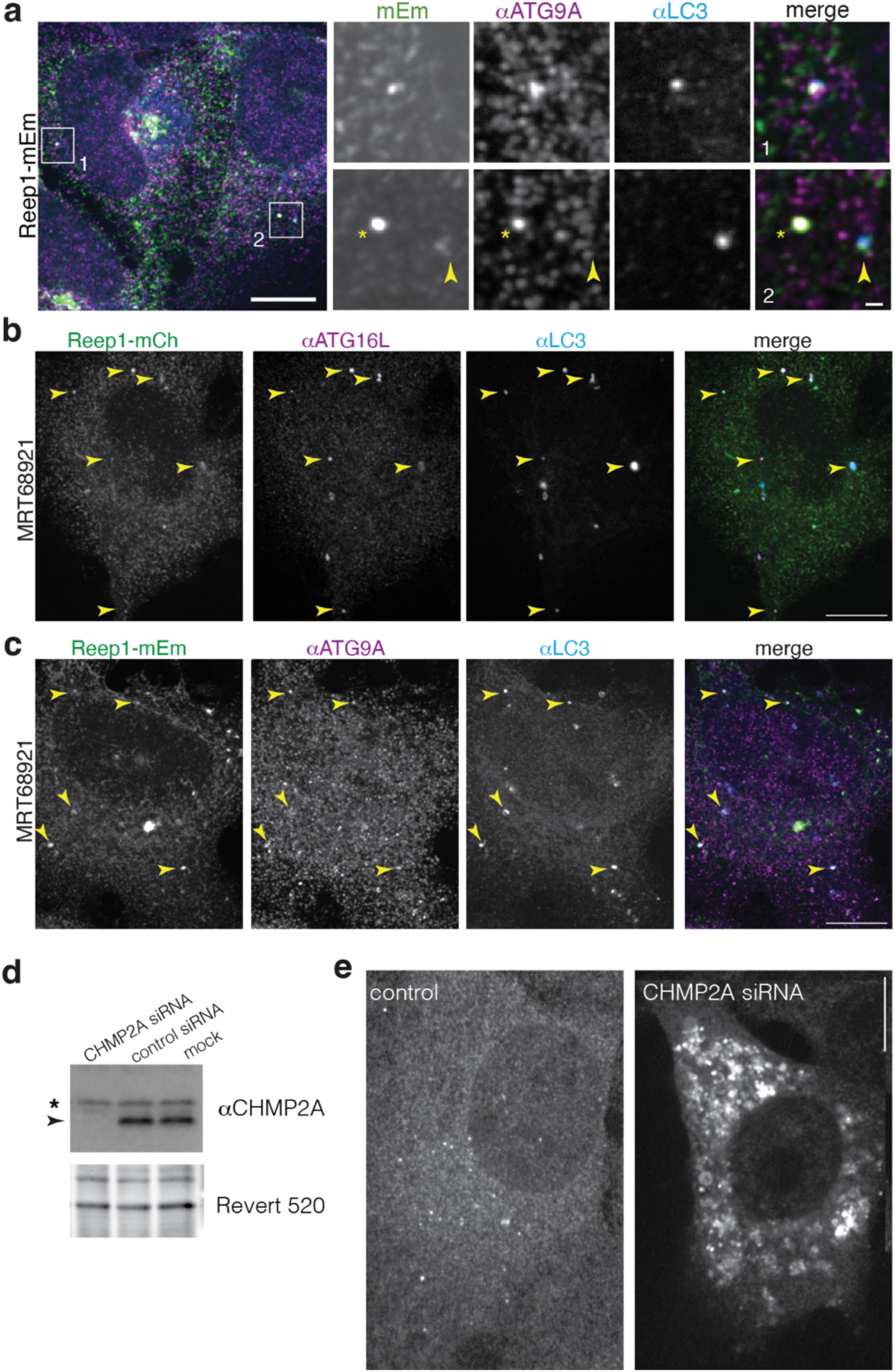
REEP1 proteins accumulate on forming autophagosomes. **a**, U2OS cells stably expressing REEP1-mEm were starved for 2 h in EBSS and immunostained for endogenous ATG9A (αATG9A) and anti-LC3 (αLC3). Right panels show magnifications of the boxed regions. Asterisk indicates a punctum containing REEP1 and ATG9A, but lacking LC3. **b**, U2OS cells stably expressing REEP1-mCh were starved in EBSS in the presence of the ULK1 inhibitor MRT38921 (1 µM) for 2h, and immunostained with anti-ATG16L (αATG16L) and LC3 (αLC3) antibodies. Arrowheads indicate examples of colocalization. **c**, As in b, but with REEP1-mEm expressing cells immunostained for endogenous ATG9A (αATG9A). **d**, Immunoblot of lysates from U2OS REEP1-mEm cells treated with CHMP2A siRNA, negative control siRNA, or mock transfected. Blots were probed with anti-CHMP2A antibodies. Arrow, CHMP2A; asterisk, non-specific band. Revert 520 stain was used as a loading control. **e**, U2OS REEP1-mEm cells treated with negative control (left panel) or CHMP2A (right panel) siRNAs were immunostained for LC3. Note the proliferation and enlargement of LC3 foci with CHMP2A depletion. Scale bars for whole cells in a, b, c, e, 10 µm; for magnification in a, 1 µm.

