## Supplemental Table 1 for "Membrane curvature-generating proteins crucial for autophagosome formation"

Supplementary Table 1. *S. pombe* strains used in this study.

| **Name** | **Genotype** |
| --- | --- |
| Sp1 | *S. pombe 972 h^-^* |
| Sp2 | *h^+^ ade6-M216 leu1-32* |
| Sp7 | *h^+^ rop1Δ::natMX6 ade6-M216 leu1-32* |
| Sp11 | *h^+^ yop1-tdTomato-natMX6 ade6-M216 leu1-32* |
| Sp19 | *h^+^ yop1-GFP-natMX ade-M216 leu1-32* |
| Sp21 | *rtn1Δ::kanMX yop1-GFP-natMX ade6 leu1-32* |
| Sp22 | *rop1Δ::kanMX6 yop1-tdTomato-natMX6 ade6-M216 leu1-32* |
| Sp25 | *rop1Δ::kanMX6 yop1-GFP-natMX ade-M216 leu1-32* |
| Sp26 | *epr1Δ::kanMX6 yop1-GFP-natMX ade-M216 leu1-32* |
| Sp27 | *atg8Δ::kanMX6 yop1-GFP-natMX ade-M216 leu1-32* |
| Sp28 | *scs2Δ::kanMX6 yop1-GFP-natMX ade-M216 leu1-32* |
| Sp32 | *h^-^ atg8Δ::natMX6 ade6-M210 leu1-32* |
| Sp33 | *h^+^ kanMX6-pnmt1-mEGFP-atg8* |
| Sp35 | *epr1Δ::natMX6 yop1-tdTomato-natMX6 ade6 leu1-32* |
| Sp36 | *atg8Δ::natMX6 yop1-tdTomato-natMX6 ade6 leu1-32* |
| Sp38 | *h^-^ rop1-5FLAG-kanMX6 ade6-M210 leu1-32* |
| Sp41 | *rop1Δ::kanMX6 kanMX6-pnmt1-mEGFP-atg8* |
| Sp43 | *h^+^ yop1Δ::kanMX ade-M216 leu1-32* |
| Sp52 | *h^-^ rop1(1-97)-SBP-kanMX6 ade6-M210 leu1-32* |
| Sp53 | *h^-^ rop1(1-123)-SBP-kanMX6 ade6-M210 leu1-32* |
| Sp69 | *h^-^ rop1(1-147)-SBP-kanMX6 ade6-M210 leu1-32* |
| Sp71 | *h^-^ kanMX6-patg8-mEGFP-atg8 ade6-M210 leu1-32* |
| Sp81 | *h^-^ rop1(W138A,V141A)-SBP-kanMX6 ade6-M210 leu1-32* |
| Sp82 | *h^+^ sec24-GFP-ura4^+^ ade6-M216 leu1-32 ura4-D18* |
| Sp82 | *h^+^ sec24-GFP-ura4 ade6-M216 leu1-32 ura4-D18* |
| Sp84 | *h^+^ anp1-GFP-ura4 ade6-216 leu1-32 ura4-D18* |
| Sp86 | *h^+^ sec72-GFP-ura4 ade6-216 leu1-32 ura4-D18* |
| Sp92 | *rop1Δ::natMX6 kanMX6-patg8-mEGFP-atg8 ade6 leu1-32* |
| Sp97 | *h^-^ pnmt1-YFP-leu1^+^ ade6 leu1-32* |
| Sp99 | *h^-^ rop1Δ::natMX6 pnmt1-YFP-leu1^+^ ade6 leu1-32* |
| Sp124 | *rop1Δ::natMX6 sec24-GFP-ura4^+^ ade6-M216 leu1-32 ura4-D18* |
| Sp124 | *rop1Δ::natMX6 sec24-GFP-ura4 ade6-M216 leu1-32 ura4-D18* |
| Sp129 | *rop1Δ::natMX6 sec72-GFP-ura4 ade6-M216 ura4-D18 leu1-32* |
| Sp138 | *rtn1-mRFP-kanMX6 ade6-M216 leu1-32* |
| Sp139 | *rop1Δ::natMX6 rtn1-mRFP-kanMX6 ade6-M216 leu1-32* |
| Sp141 | *sey1Δ::natMX rtn1-mRFP-kanMX6 ade6-M216 leu1-32* |
| Sp143 | *yop1-mRFP-kanMX6 ade6-M216 leu1-32* |
| Sp144 | *rop1Δ::natMX6 yop1-mRFP-kanMX6 ade6-M216 leu1-32* |
| Sp146 | *sey1Δ::natMX yop1-mRFP-kanMX6 ade6-M216 leu1-32* |
| Sp149 | *h^+^ fsc1Δ::hphMX ade6-M216 leu1-32 ura4-D18* |
| Sp151 | *h^-^ fsc1Δ::hphMX rop1Δ::natMX6 ade6-M216 leu1-32 ura4-D18* |
| Sp152 | *h^-^ cpy1-mNeonGreen-hphMX6 ade6-M210 leu1-32 ura4-D18* |
| Sp157 | *rop1-mNeonGreen-natMX6 yop1-mRFP-kanMX6 ade6-M216 leu1-32* |
| Sp162 | *kanMX6-pnmt1-mEGFP-atg8 fsc1Δ::hphMX ade6-M216 leu1-32 ura4-D18* |
| Sp163 | *kanMX6-pnmt1-mEGFP-atg8 fsc1Δ::hphMX rop1Δ::natMX6 ade6-M216 leu1-32 ura4-D18* |
| Sp170 | *cpy1-mNeonGreen-hphMX6 rop1Δ::natMX6 ade6-M210 leu1-32 ura4-D18* |
| Sp175 | *h^+^ rop1(Δ133-147)-SBP-kanMX6 ade6-M216 leu1-32* |
| Sp197 | *h^+^ vps24-mEGFP-hphMX6 ade6-M210 leu1-32 ura4-D18* |
| Sp198 | *h^+^ vps28-mEGFP-hphMX6 ade6-M210 leu1-32 ura4-D18* |
| Sp208 | *sfb3-mCherry-hphMX6 ade6 leu1-32* |
| Sp209 | *rop1Δ::natMX6 sfb3-mCherry-hphMX6 ade6 leu1-32* |
| Sp228 | *h^-^ rop1(Δ97-105)-SBP-kanMX6 ade6-M210 leu1-32* |
| Sp229 | *h^-^ rop1(Δ106-114)-SBP-kanMX6 ade6-M210 leu1-32* |
| Sp230 | *h^-^ rop1(Δ166-181)-SBP-kanMX6 ade6-M210 leu1-32* |
| Sp245 | *tom20-mCherry-hphMX6 ade6 leu1-32* |
| Sp246 | *rop1Δ::natMX6 tom20-mCherry-hphMX6 ade6 leu1-32* |
| Sp248 | *atg8Δ::kanMX6 tom20-mCherry-hphMX6 ade6 leu1-32* |
| Sp265 | *h^+^ tdh1-mCherry-hphMX6 ade6 leu1-32* |
| Sp266 | *rop1Δ::natMX6 tdh1-mCherry-hphMX6 ade6 leu1-32* |
| Sp267 | *atg8Δ::kanMX6 tdh1-mCherry-hphMX6 ade6 leu1-32* |
| Sp269 | *h^-^ pyk1-mCherry-hphMX6* |
| Sp270 | *h^-^ pgk1-mCherry-hphMX6* |
| Sp272 | *h^-^ hsc1-GFP-hphMX ade6-M210 leu1-32* |
| Sp274 | *rop1Δ::natMX6* |
| Sp275 | *atg8Δ::natMX6* |
| Sp276 | *yop1Δ::kanMX* |
| Sp278 | *pgk1-mCherry-hphMX6 rop1Δ::natMX6* |
| Sp279 | *pgk1-mCherry-hphMX6 atg8Δ::kanMX6* |
| Sp280 | *hsc1-GFP-hphMX rop1Δ::natMX6 ade6-M210 leu1-32* |
| Sp281 | *hsc1-GFP-hphMX atg8Δ::kanMX6 ade6-M210 leu1-32* |
| Sp282 | *h^-^ pex11-mCherry-hphMX6* |
| Sp285 | *pyk1-mCherry-hphMX6 rop1Δ::natMX6* |
| Sp286 | *pyk1-mCherry-hphMX6 atg8Δ::kanMX6* |
| Sp293 | *kanMX6-p41nmt1-mCherry-atg8 kanMX6-p81nmt1-rop1-mNeonGreen-natMX6 ade6 leu1-32* |
| Sp294 | *rop1Δ::natMX6 pex11-mCherry-hphMX6* |
| Sp295 | *atg8Δ::kanMX6 pex11-mCherry-hphMX6 ade6* |
| Sp298 | *h^-^ rop1Δ::prop1-atg40-5FLAG-kanMX6 ade6-M210 leu1-32* |
| Sp336 | *atg2-GBP-mcherry-kanMX6 yop1-GFP-natMX leu1-32* |
| Sp356 | *h^+^ sec63-GFP-natMX* |
| Sp358 | *atg2-tdTomato-hphMX6 kanMX6-patg8-mEGFP-atg8 ade6-M216 leu1-32* |
| Sp359 | *rop1Δ::natMX6 atg2-tdTomato-hphMX6 kanMX6-patg8-mEGFP-atg8 ade6-M216 leu1-32* |
| Sp364 | *rop1Δ::hphMX atg2-GBP-mcherry-kanMX6 yop1-GFP-natMX* |
| Sp370 | *rop1Δ::natMX6 anp1-GFP-ura4 ade6-216 leu1-32 ura4-D18* |
| Sp378 | *rop1Δ::kanMX6 sec63-GFP-natMX* |
| Sp380 | *rop1Δ::hphMX atg2-GBP-mcherry-kanMX6 sec63-GFP-natMX* |
| Sp391 | *rop1Δ::hphMX yop1-GFP-natMX atg18a-GBP-mcherry-kanMX6 leu1-32* |
| Sp392 | *rop1Δ::hphMX yop1-GFP-natMX atg18b-GBP-mcherry-kanMX6 ade6-M216* |
| Sp394 | *rop1Δ::natMX6 vps24-mEGFP-hphMX6 ade6-M210 leu1-32 ura4-D18* |
| Sp395 | *rop1Δ::natMX6 vps28-mEGFP-hphMX6 ade6-M210 leu1-32 ura4-D18* |
| Sp404 | *rop1Δ::hphMX atg2-GBP-mcherry-kanMX6 yop1(d132-189)-GFP-natMX* |
| Sp405 | *rop1Δ::hphMX atg2-GBP-mcherry-kanMX6 yop1(d143-189)-GFP-natMX* |
| Sp411 | *h^+^ zhf1-mNeonGreen-kanMX6 ade6-M216 leu1-32* |
| Sp412 | *atg1-tdTomato-hphMX6 kanMX6-patg8-mEGFP-atg8* |
| Sp413 | *rop1Δ::natMX6 atg1-tdTomato-hphMX6 kanMX6-patg8-mEGFP-atg8* |
| Sp418 | *atg9-tdTomato-hphMX6 kanMX6-patg8-mEGFP-atg8 ade6-M216 leu1-32* |
| Sp424 | *rop1Δ::natMX6 atg9-tdTomato-hphMX6 kanMX6-patg8-mEGFP-atg8 leu1-32* |
| Sp434 | *rop1Δ::natMX6 zhf1-mNeonGreen-kanMX6 ade6-M216 leu1-32* |
| Sp445 | *h^-^ kanMX6-patg8-mEGFP-atg8 atg5-mcherry-hphMX6 ade6-M210 leu1-32* |
| Sp446 | *h^-^ rop1Δ::natMX6 kanMX6-patg8-mEGFP-atg8 atg5-mcherry-hphMX6 ade6 leu1-32* |
| Sp454 | *pyop1-rop1-mcherry-leu1 yop1-GFP-natMX ade6 leu1-32* |
| Sp471 | *h^+^ rop1-APEX2-flag-natMX6 ade6-M216 leu1-32* |
| Sp473 | *h^+^ atg2-APEX2-flag-natMX6 ade6-M216 leu1-32* |
| Sp474 | *h^+^ atg2-APEX2-flag-natMX6 ade6-M216 leu1-32* |
