## Supplemental Table 2 for "Membrane curvature-generating proteins crucial for autophagosome formation"

Supplementary Table 2. Plasmids used to express proteins in *E. coli* and *S. pombe*.

| Name | | Note |
| --- | --- | --- |
| NWP385 | pET21b-sjRop1-3C-SBP | Express sjRop1 in *E. coli* |
| NWP390 | pET21b- sjRop1(Δ126-170)-3C-SBP | Express sjRop1 mutant in *E. coli* |
| NWP391 | pET21b- sjRop1(Δ107-120)-3C-SBP | Express sjRop1 mutant in *E. coli* |
| NWP395 | pET28-hsREEP1-TEV-SBP | Express human REEP1 in *E. coli* |
| NWP485 | pET28-hsREEP1(L107P)-TEV-SBP | Express human REEP1 mutant in *E. coli* |
| NWP486 | pET28-hsREEP1(Δ102-139)-TEV-SBP | Express human REEP1 mutant in *E. coli* |
| NWP481 | pET21b- sjRop1(V16Bpa)-3C-SBP | V16 codon was mutated to TAG for incorporation of Bpa at this position; Amber codon incorporated at other positions of TM1 are available for request |
| NWP412 | pET21b-His10-TEV-scAtg8(1-116) | Express scAtg8 amino acid 1-116 in *E. coli* |
| NWP451 | pGEX-6p-spAtg8 | Express spAtg8 in *E. coli* |
| NWP410 | pET28-scAtg40-TEV-SBP | Express scAtg40 in *E. coli* |
| NWP419 | pET28-scAtg40(Y242A, M245A)-TEV-SBP | Express scAtg40 LIR mutant in *E. coli* |
| NWP413 | pET21b-sjRop1(W144A, I147A)-3C-SBP | Express sjRop1 LIR mutant in *E. coli* |
| NWP435 | pJK148-*nmt1*-EYFP | To integrate and express EYFP (fused with HHGNSGPPPPGAFPHPLEGGDPPVAT at N-terminus) at *leu1* locus in *S. pombe* |
| NWP436 | pFA6a-kanMX6-*patg8*-mEGFP | To express mEGFP N-terminally tagged spAtg8 at its genomic locus |
| NWP463 | Topo-*prop1*-Rop1-5xFLAG-kanMX6 | To express Rop1 tagged with 5xFLAG at its genomic locus; deletion or point mutations affecting APH and predicted LIR motif of Rop1 are available for request |
| NWP504 | Topo-*prop1*-Atg40-5FLAG-kanMX6 | To express Atg40 tagged with 5xFLAG at Rop1’s genomic locus |
